# Local adaptation under gene flow: Recombination, conditional neutrality and genetic trade-offs shape genomic patterns in *Arabidopsis lyrata*

**DOI:** 10.1101/374900

**Authors:** Tuomas Hämälä, Outi Savolainen

## Abstract

Short-scale local adaptation is a complex process involving selection, migration and drift. The expected effects on the genome are well grounded in theory, but examining these on an empirical level has proven difficult, as it requires information about local selection, demographic history and recombination rate variation. Here, we use locally adapted and phenotypically differentiated *Arabidopsis lyrata* populations from two altitudinal gradients in Norway to test these expectations at the whole-genome level. Demography modelling indicates that populations within the gradients diverged less than 2000 years ago and that the sites are connected by gene flow. The gene flow estimates are, however, highly asymmetric with migration from high to low altitudes being several times more frequent than *vice versa*. To detect signatures of selection for local adaptation, we estimate patterns of lineage specific differentiation among these populations. Theory predicts that gene flow leads to concentration of adaptive loci in areas of low recombination; a pattern we observe in both lowland-alpine comparisons. Although most selected loci display patterns of conditional neutrality, we found indications of genetic trade-offs, with one locus particularly showing high divergence and signs of selection in both populations. Our results further suggest that resistance to solar radiation is an important adaptation to alpine environments, while vegetative growth and bacterial defense are indicated as selected traits in the lowland habitats. These results provide insights into genetic architectures and evolutionary processes driving local adaptation under gene flow. We also contribute to understanding of traits and biological processes underlying alpine adaptation in northern latitudes.

## Introduction

Population genetics theory by Haldane (Haldane 1930) and Wright (Wright 1931) laid the basis for predicting how selection and drift influence adaptive variation in the presence of gene flow. More recently, theoretical studies have shown that the interplay between these factors can lead to characteristic changes in genetic architectures underlying local adaptation (Griswold 2006; Kirkpatrick and Barton 2006; Bürger and Akerman 2011; Yeaman and Whitlock 2011; Akerman and Bürger 2014). An important prediction is the shift towards fewer large effect loci that are clustered in areas of reduced recombination, as correlated sites under strong selection are less easily swamped by gene flow (Lenormand and Otto 2000; Lenormand 2002; Kirkpatrick and Barton 2006; Bürger and Akerman 2011; Yeaman and Whitlock 2011; Akerman and Bürger 2014). The generality of this prediction has been shown with a continent-island model (single population adapting to a new environment under maladaptive migration) (Bürger and Akerman 2011; Akerman and Bürger 2014), as well as a framework where two connected populations experience selection towards distinct optima (Yeaman and Whitlock 2011; Akerman and Bürger 2014). The adaptive loci are also expected to show antagonistic pleiotropy, wherein a single locus is kept polymorphic by differential selection to contrasting environments, because under the alternative scenario of conditional neutrality (allele is selected for or against only in one environment, while being neutral in the other), the overall beneficial allele will be fixed at all populations over time (Yeaman and Whitlock 2011; Savolainen et al. 2013; Akerman and Bürger 2014; Wadgymar et al. 2017; Yoder and Tiffin 2017). The footprint of positive and negative selection on linked variants [i.e. genetic hitchhiking (Maynard Smith and Haigh 1974) and background selection (Charlesworth et al. 1993), respectively] can be intensified by gene flow (Laurent et al. 2016; Pfeifer et al. 2018; Wang et al. 2018), but as migration also hinders adaptive differentiation, local adaptation under gene flow is restricted to environments where sharp differences in spatially variation selection can exist. For example, plant populations growing on toxic soils, either naturally occurring or contaminated by mine tailings, have exhibited such adaptation (Antonovics and Bradshaw 1970; Sambatti and Rice 2006; Arnold et al. 2016; Aeschbacher et al. 2017).

In the present study, we examine the genetic basis of local adaptation among lowland and alpine populations of a self-incompatible perennial plant, *Arabidopsis lyrata*. In mountainous environments, abiotic factors such as temperature and solar radiation can differ drastically among closely adjacent areas, producing steep environment gradients (Gonzalo-Turpin and Hazard 2009; Fischer et al. 2013; Kubota et al. 2015; Günther et al. 2016), not unlike those found on toxic soils. Indeed, we have recently demonstrated that *A. lyrata* individuals from different altitudes exhibit population specific phenotypes when grown at native common garden sites in Norway and in a novel habitat in Finland (Hämälä et al. 2018). Using whole-genome based demography modelling and a reciprocal transplant experiment, we further showed that despite evidence of gene flow, local alpine and lowland populations have highest fitness at their home sites, satisfying the local vs. foreign criterion of local adaptation (Kawecki and Ebert 2004). This study system and the combined knowledge of local adaptation and demographic history, as well as genome-wide recombination information from a high density linkage map (Hämälä et al. 2017), provides us a rare opportunity to study genomic patterns of adaptive differentiation under gene flow (Savolainen et al. 2013).

Here, we extend our previous work to the genome level by examining variation potentially underlying the local adaptation. If the alpine adaptation evolved after recent colonization from the lowland sites, the high-altitude populations are expected to show lower genetic diversities and stronger population size contractions than the low altitude populations. The ongoing gene flow then leads us to look for evidence of antagonistic pleiotropy (loci showing high differentiation and signs of selection in both populations) and loci where reduced recombination contributes to maintaining polymorphisms. Furthermore, we expect to identify genes and biological processes underlying the lowland and alpine adaptation. We search for signs of selection by employing a measure for lineage specific differentiation, the population branch statistic (PBS) (Yi et al. 2010), on a set of whole-genome sequences. Unlike traditional differentiation measures, PBS can distinguish the selected lineage by combining information from two closely related populations and an outgroup. As this method searches for allele frequency differences among populations, it is more likely to detect large effect loci instead of small effect ones underlying truly quantitative traits, because the latter signal may largely arise from linkage disequilibrium between populations (Latta 1998; Berg and Coop 2014). We then explore the potential effects of selection with coalescent and forward simulations, by utilizing parameter estimates available for these populations.

We specifically address the following questions: What do diversity patterns tell us about the demographic history of these populations, and do we find footprints of gene flow on this variation? Do we find more selected sites in areas of reduced recombination in populations that receive more migrants? Are the adaptive loci more clustered under higher gene flow? Is the genetic architecture dominated by conditional neutrality or do we find loci exhibiting variation consistent with antagonistic pleiotropy? And to what phenotypes and biological processes are the outliers associated with and do we find evidence of adaptive convergence between the two alpine areas?

## Results

We used whole-genome data from 47 resequenced *A. lyrata* individuals. Most of the samples were collected from four populations growing in two alpine areas in Norway, each represented by a high and low group (Jotunheimen and Trollheimen; Fig 1): Lom, Jotunheimen (J1, 300 m.a.s.l.; *n* = 9), Spiterstulen, Jotunheimen (J3, 1100 m.a.s.l; *n* = 12), Sunndalsøra, Trollheimen (T1, 10 m.a.s.l; *n* = 5) and Nedre Kamtjern, Trollheimen (T4, 1360 m.a.s.l; *n* = 9). The abbreviations come from Hämälä et al. (2018). For some of the analysis, we also included samples from Germany (GER; *n* = 6) and Sweden (SWE; *n* = 6) as comparison groups (Table S1). Data from the T4 population represent new collections for the current study, whereas other samples came from two earlier studies (Mattila et al. 2017; Hämälä et al. 2018).

**Figure 1.**
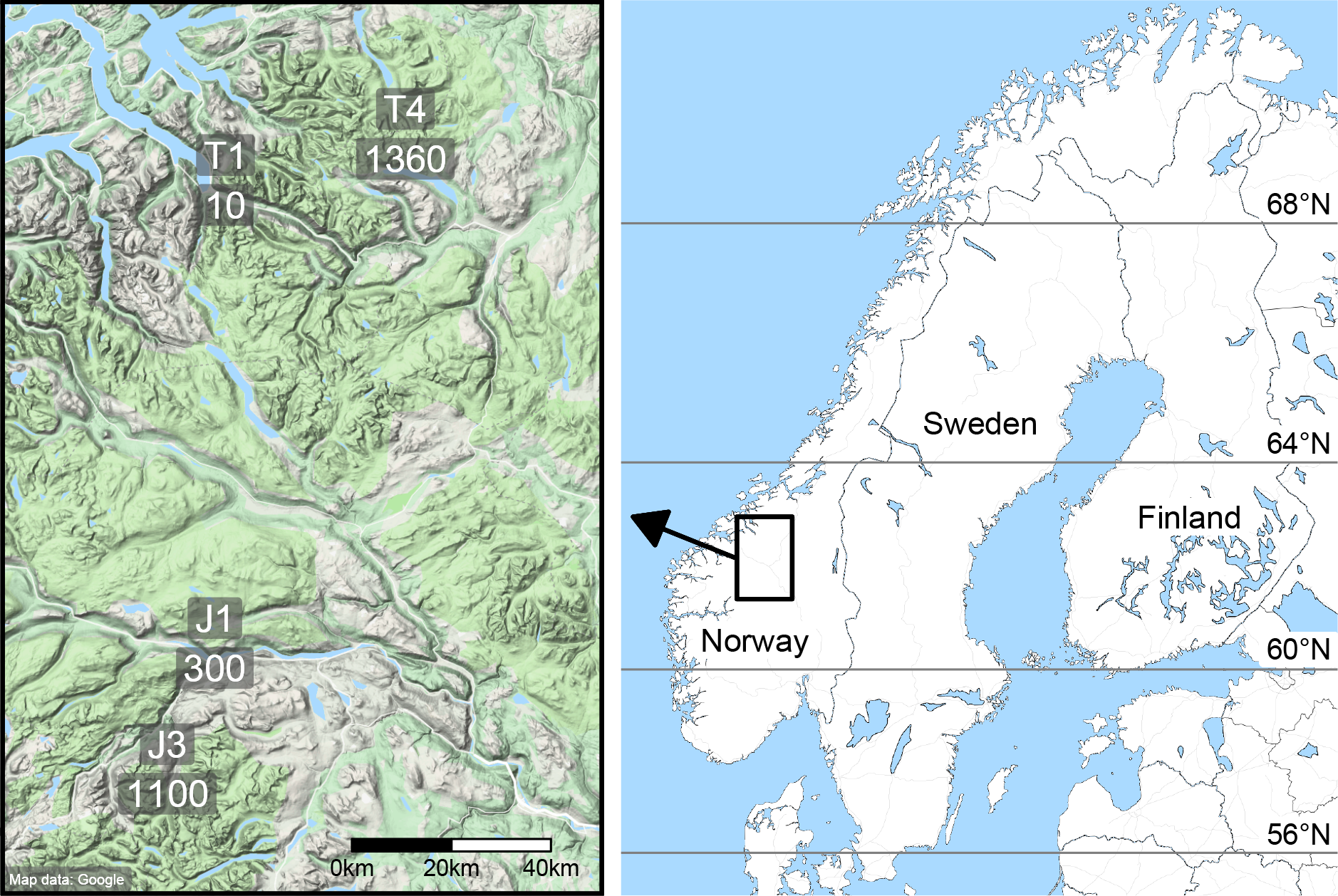
Locations and altitudes of the *A. lyrata* growing sites.

**Figure 2.**
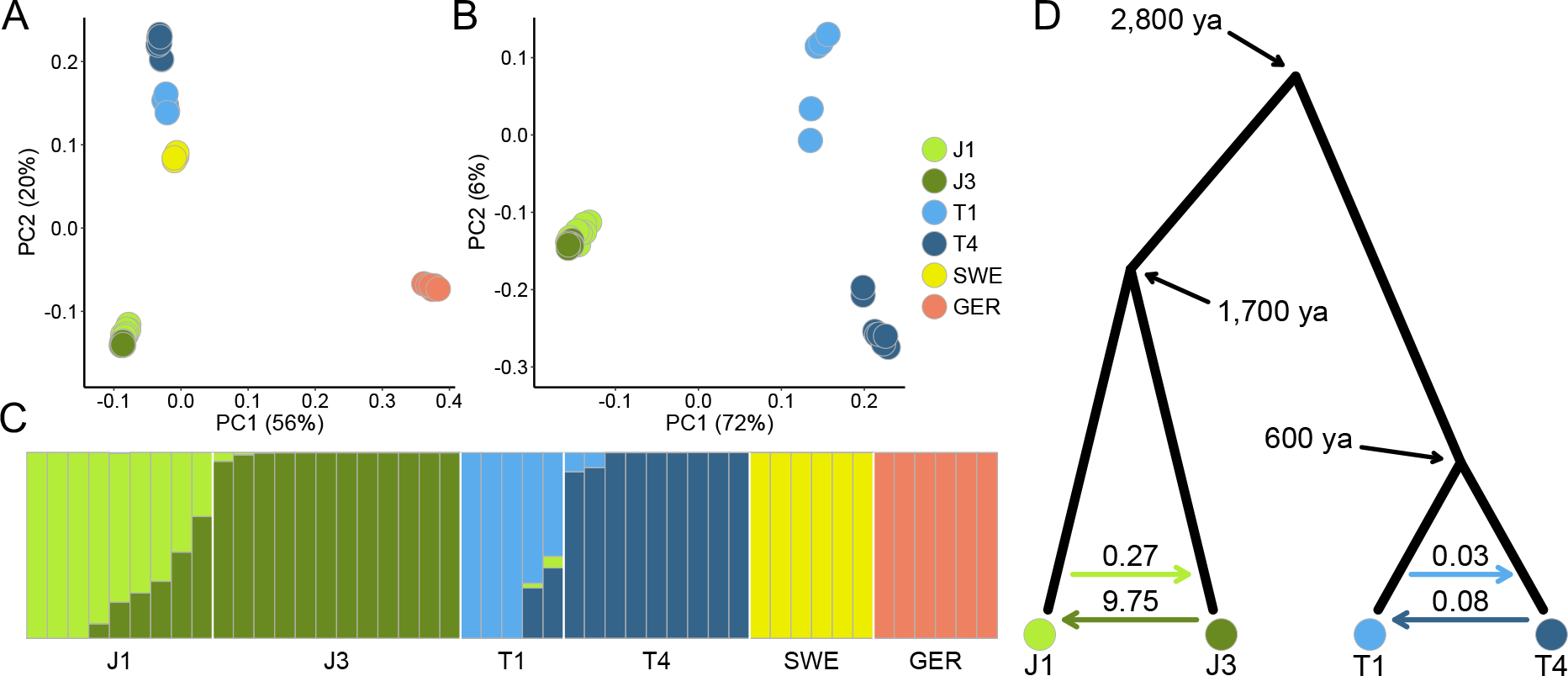
Visualization of the population structure and demographic history. Besides the Norwegian populations (J1, J3, T1, T4), individuals from Sweden (SWE) and Germany (GER) are included for comparison. **A:** Genetic variation along the first two eigenvectors of a principal component analysis (PCA). All six populations included. **B:** PCA with only the Norwegian populations included. The percentage of variance explained by the principal components are shown in brackets. **C:** Estimated admixture proportions for the best supported number of ancestral populations (K = 6). **D:** Estimated divergence times and migration rates between the Norwegian populations. Times are in years, while assuming a generation time of two years. Estimates above the colored arrows indicate population migration rates (4*N*_*e*_*m*).

### Alpine populations harbour lower genetic diversities than lowland populations

Within each area, the high-altitude populations had lower synonymous nucleotide diversities than the low-altitude populations (*p* < 2.2×10^−16^; Fig S1). Synonymous Tajima’s *D* estimates were also more highly positive in the high-altitude populations (*p* < 2.2×10^−16^; Fig S2), suggesting stronger population size contractions in their history. Differentiation levels, as estimated with *F*_ST_, were lower between the neighbouring lowland and alpine populations than between other comparison pairs (Table S2). The Jotunheimen populations were less differentiated from each other (J1-J3 *F*_ST_ = 0.09) than populations from Trollheimen (T1-T4 *F*_ST_ = 0.17).

We then conducted a principal component analysis (PCA) and an admixture analysis to further evaluate relationships between the studied individuals. PCA showed distinct divisions, with individuals clustering according to populations and populations clustering according to geographical location (Fig 1A). Consistent with the *F*_ST_ estimates, the Trollheimen populations were more clearly separated from each other than populations from Jotunheimen. In fact, in a model with only the focal populations included, variation along the first two principal components did not separate the J1 and J3 populations (separation happens at PC4; Fig S3). The admixture patterns were in most part concordant with the PCA results. In general, individuals from the low-altitude populations J1 and T1 had higher admixture proportions (from their respective high-altitude neighbours) than individuals from the high-altitude populations J3 and T4 (from their low-altitude neighbours) (Fig 1C).

### Demography analysis reveals recent divergence and asymmetric gene flow

To quantify the levels of gene flow, as well as to have an estimate of divergence times and effective population sizes, we conducted site frequency spectra based demography simulations in fastsimcoal2 (Excoffier et al. 2013). Simulations involving the Jotunheimen populations J1 and J3 were done as part of an earlier study (Hämälä et al. 2018) and here we add parameter estimates for the Trollheimen populations T1 and T4. Maximum likelihood estimates (MLE) from the best-supported models (for model selection, see Table S3) indicated more recent divergence between T1 and T4 (307 generations ago) populations then between J1 and J3 (866 generations ago) populations. The Trollheimen groups also had lower effective population size estimates (*N*_*e*_: T1 = 1862; T4 = 779) than the Jotunheimen groups (*N*_*e*_: J1 = 3370; J3 = 4295). Although the levels of gene flow were clearly higher among the Jotunheimen populations, population migration rates (4*N*_*e*_*m*) were heavily biased towards the low altitude populations in both alpine areas (~36× higher from J3 to J1 than from J1 to J3, and ~2.4× higher from T4 to T1 than from T1 to T4) (Fig 1D). As *A. lyrata* is insect pollinated, the asymmetry in gene flow could result from higher seed dispersal or pollinator movement from high- to low- altitudes, but because allele frequencies reflect events over extended periods of time, we cannot rule out other modes of migration that may have been in effect during colonization of the alpine areas. For all MLEs and their confidence intervals, see Table S4.

### Genome-wide patterns of population specific selection

We studied population specific differentiation to examine the effects of gene flow on neutral and selected variants and to ascertain important biological processes at each lowland and alpine habitat. To this end, we used population branch statistic (PBS) (Yi et al. 2010) to distinguish loci that have been under directional selection after the adjacent low- and high-altitude populations diverged. Selection patterns between the neighbouring populations were examined by calculating PBS estimates in 50 SNP non-overlapping sliding windows for population trios J1-J3-GER and T1-T4-GER. Using neutral data simulated under the best supported demography models, we first generated ~50,000 PBS samples for each population and used the distributions to find limits to neutral variation in the observed data. The simulations predicted lower median estimates and more tightly centred distributions in the low-altitude populations than in the high-altitude populations (Fig 3A). Overall, the observed estimates corresponded well with the simulated data. Within both areas, the low-altitude populations had lower median estimates than the high-altitude population and the distributions were less dispersed around the median (Fig 3A). The observed distributions did, however, have slightly longer lower and upper tails than the neutral ones, suggesting balancing and directional selection, respectively. Significant deviations from the neutral expectations were determined by comparing the observed estimates against quantiles of the simulated distributions. Our outlier detection model detected fewer selected loci in the low-altitude populations than in the high-altitude populations (Table 1). Although all outlier windows tended to localize in areas with lower than average recombination rates (genome-wide average = 3.7×10^−8^ crossing-overs per base pair), this trend was more pronounced in the low-altitude populations (*p* < 0.006; Fig S4) (Table 1). As the theory further predicts that gene flow leads to clustering of the adaptive loci, we measured for each outlier window the distance to the closest adjacent outlier window. This measure, as opposed to using all between-window distances, is less affected by distances between the potential clusters. In both altitude comparisons, outliers in the low-altitude populations were located marginally closer to each other than in the high-altitude populations (*p* < 0.02; Fig S5) (Table 1), but there was no significant correlation between distance and recombination rate (Pearson *r* = 0.022, *p* = 0.48; Fig S6). To explore additional factors that might explain the occurrence of the outliers in low recombination regions, we examined variation in gene densities and mutation rates across the *A. lyrata* genome. We obtained a proxy for mutation rate by estimating the rate of synonymous substitutions (*d*_S_) between *A. lyrata* and *A. thaliana*. A positive correlation was observed between recombination rate and gene density (Pearson *r* = 0.159, *p* < 2.2×10^−16^; Fig S7) and recombination rate and *d*_S_ (Pearson *r* = 0.109, *p* < 2.2×10^−16^; Fig S8), but neither gene density (*p* > 0.09; Fig S9) nor *d*_S_ (*p* > 0.24; Fig S10) showed significant differences in outlier areas between the low- and high-altitude populations. Lastly, to assess differences in recombination rate estimates between populations while controlling for gene density and mutation rate variation, we fitted the following linear model to our Jotunheimen and Trollheimen outlier sets: recombination rate = gene density + *d*_S_ + population. For both data sets, likelihood-ratio tests comparing the fit of a full model to a reduced model with the population variable removed indicated significant differences between the low- and high-altitude populations (*p* < 0.05). Thus, the fact that outliers in the lowland populations are found in areas with lower recombination rates is due to recombination itself, not correlated genomic characteristics.

**Figure 3.**
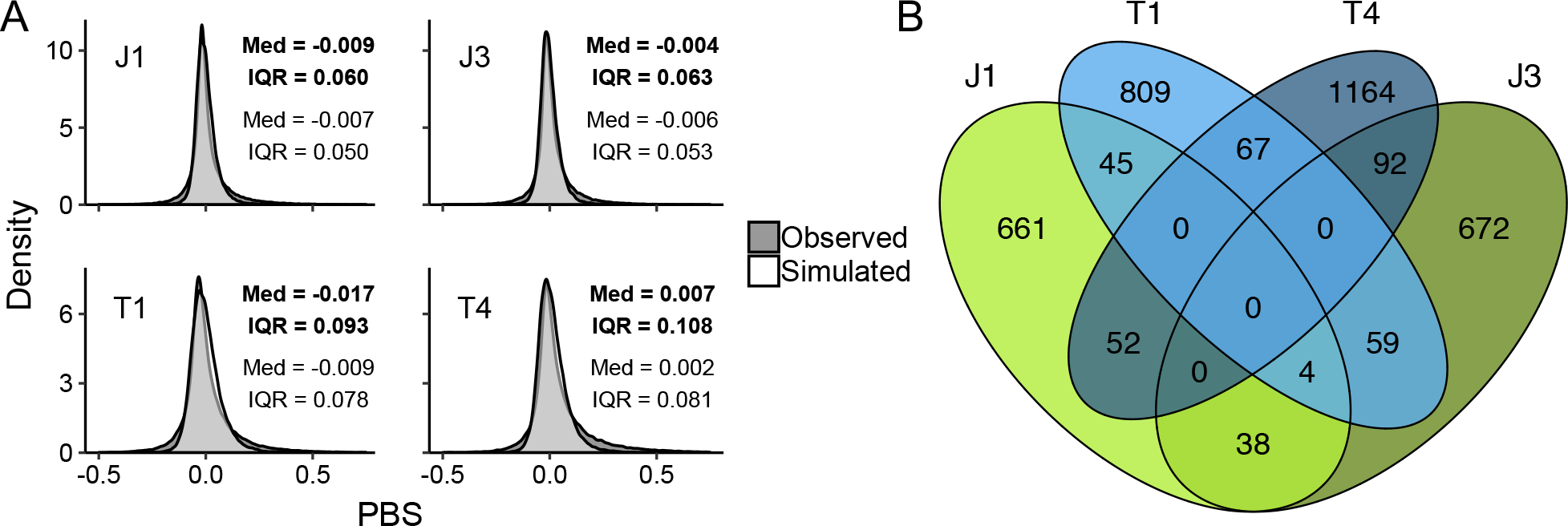
Identification of putatively adaptive loci. **A:** Observed population branch statistic (PBS) distributions compared against simulated neutral samples. Median estimates and interquartile ranges are marked for the observed (bold) and simulated (plain) distributions. **B:** Number of annotated genes found within 5 Kb of the significant (*q* < 0.05) PBS outlier windows.

**Table 1.** Number of outlier windows, within outlier window recombination rates and between outlier window distances for each population.

| | #outliers | | Recombination rate ($\times 10^{-8}$) | | Distance (Kb) | |
| --- | --- | --- | --- | --- | --- | --- |
|  | 50-SNP | 1-SNP | 50-SNP | 150-SNP | 50-SNP | 150-SNP |
| J1 | 896<br>(1.1%) | 2.14<br>(2.11 – 2.15) | 2.03<br>(1.91 – 2.18) | 2.04<br>(1.79 – 2.28) | 3.18<br>(2.69 – 3.67) | 11.69<br>(7.63 – 14.38) |
| J3 | 948<br>(1.2%) | 2.22<br>(2.20 – 2.25) | 2.38<br>(2.22 – 2.70) | 2.36<br>(2.13 – 2.68) | 3.80<br>(3.18 – 4.62) | 12.52<br>(9.00 – 15.21) |
| T1 | 966<br>(1.3%) | 2.08<br>(2.04 – 2.10) | 2.06<br>(1.83 – 2.30) | 1.97<br>(1.76 – 2.22) | 3.91<br>(2.99 – 4.38) | 13.12<br>(8.38 – 15.98) |
| T4 | 1160<br>(1.6%) | 2.14<br>(2.14 – 2.15) | 2.46<br>(2.36 – 2.74) | 2.55<br>(2.33 – 2.70) | 3.96<br>(3.34 – 4.47) | 14.44<br>(11.99 – 17.61) |

Median estimates and their 95% CIs are shown in different SNP-based window sizes. For the 1- and 150-SNP windows, we assumed the same percentage of outliers as estimated for the 50-SNP windows. Median genome-wide recombination rate = $3.7 \times 10^{-8}$ .

### Evidence of allelic trade-off at *XRN2* locus

Most selection outliers were found only in a single population (Dataset S1), likely caused by selection acting only on a single population (conditional neutrality). Some loci were, however, shared between the neighboring low- and high-altitude populations, suggesting possible antagonistic pleiotropy: 42 out of 1668 genes that were found within 5 Kb of an outlier window were shared between J1 and J3 (based on 10,000 permutated data sets, the expected number of shared outliers is 22; *p* = 0.0003), while 67 out of 2359 were shared between T1 and T4 (43 expected; *p* = 0.0007). (Fig 3B). One locus, *XRN2*, particularly stood out as one of the top outliers in T1 and T4, caused by higher than neutral differentiation between all three population comparisons (i.e. T1-T4, T1-GER and T4-GER). To examine whether the observed pattern can result from directional selection favoring different alleles in different populations (antagonistic pleiotropy), we examined allele frequencies, as well as pairwise nucleotide diversities (*π*) and Tajima’s *D* estimates around the gene. Reduced diversity surrounding the selected site is a classical signal of a rapid selective sweep (Maynard Smith and Haigh 1974), whereas negative Tajima’s *D* indicates skews in the site frequency spectrum due to excess of rare variants – a pattern also consistent with a selective sweep (Braverman et al. 1995). Allele frequencies showed clear evidence of differentiation, with T1 fixed for one allele and T4 nearly fixed for the alternative allele. Both populations also had reduced nucleotide diversities and negative Tajima’s *D* estimates in an ~8 Kb area around the gene, suggesting directional selection in each lineage (Fig 4). To evaluate the likelihood of the observed patterns under our estimated demography parameters, we used forward genetic models in SLiM 2 (Haller and Messer 2017) to simulate nucleotide diversities and Tajima’s *D* estimates under different selection scenarios. We ran the simulation 10 × *N*_*e*_ generations to approach mutation-drift balance and introduced the sweep mutation with beneficial selection coefficient *α*_b_ = 4*N*_*e*_*s* = 100, 1000 and 10,000. To simulate the effects of a genetic trade-off, deleterious alleles with selection coefficient *α*_d_ = ½ × – *α*_b_ migrated into a population with rate 4*N*_*e*_*m* = 0.083 or 0.034. We then recorded the time in number of generations until *π* and *D* dropped below the initial neutral estimate. The simulations were conducted as single-origin hard sweeps and as multiple-origin soft sweeps. For the soft-sweeps, we assumed that 5% of the corresponding population carried the adaptive allele before the start of selection. Each simulation was repeated 500 times and the median value retained. These simplified models show that under a hard sweep with strong trade-off (*α*_b_ > 100 and *α*_d_ < –50), *π* and *D* could respond to selection in both populations within the estimated time frame. The same, however, was not true for lower selection coefficients or multiple-origin soft sweeps in general (Fig 5).

**Figure 4.**
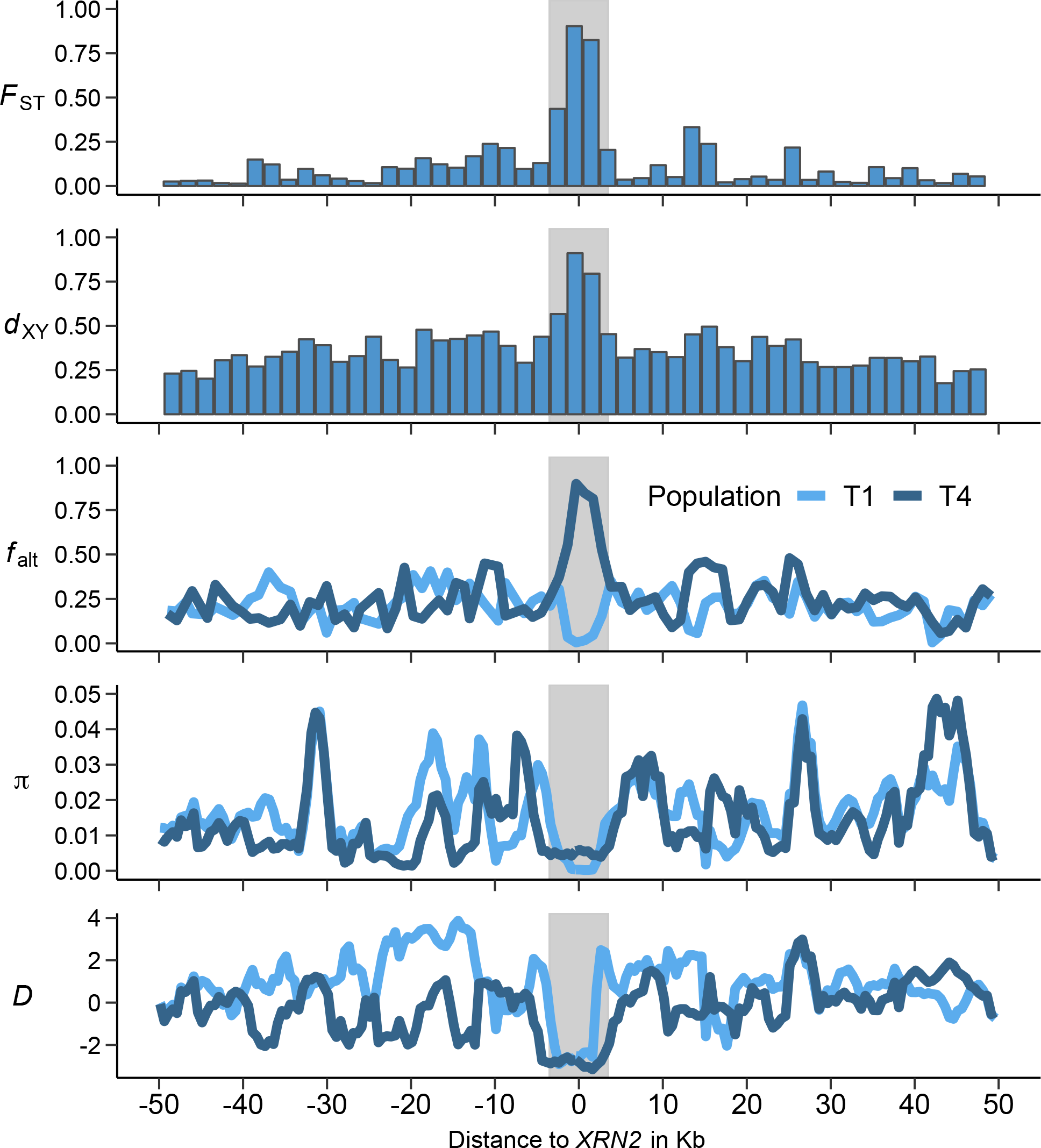
Evidence of opposing selective sweeps between T1 and T4 populations at candidate loci *XRN2*. *F*_ST_ gives the relative and *d*_XY_ the absolute allele frequency differentiation at variable sites. *f*_alt_ shows frequencies of the alternate non-reference SNP alleles. Nucleotide diversity *π* and Tajima’s *D* indicate a loss of heterozygosity and an excess of rare variants, respectively. Shading marks the coding area of the gene.

**Figure 5.**
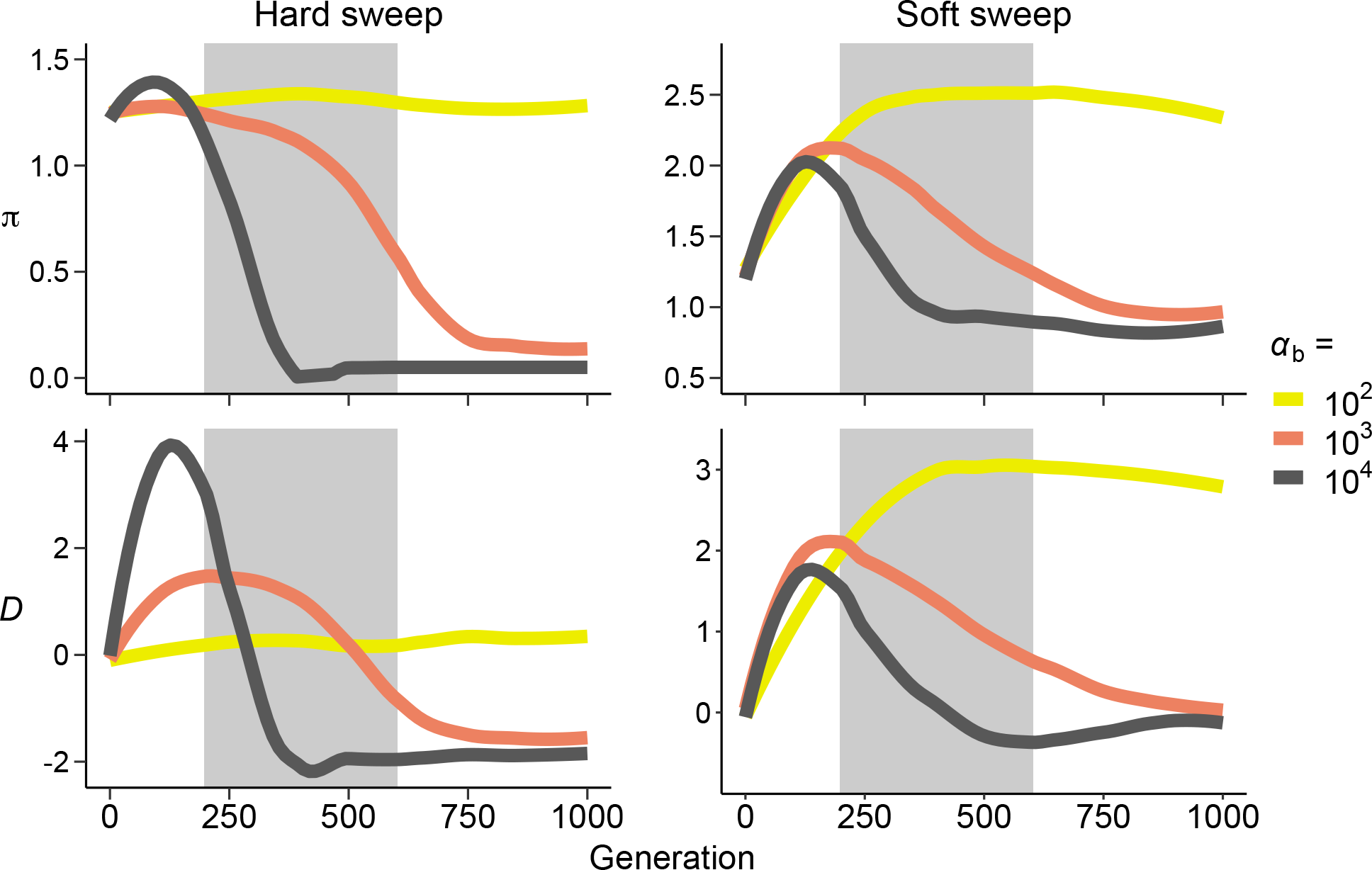
Insights into selection acting on *XRN2*. Nucleotide diversity *π* (×10^−3^) and Tajima’s *D* were simulated for 8 Kb area with parameters corresponding to T1 population. The simulations were ran as single-origin hard sweeps and as multiple-origin soft sweeps. Shown are median estimates from 500 simulations. Shaded area marks the 95% confidence intervals for the estimated divergence time between T1 and T4 populations. For T4 population, see Fig S11.

### Biological processes show adaptive convergence

Most genes that localized within 5 Kb of the outlier windows were population specific (Dataset S1). Among the significant outlier loci, 49 out of 1785 genes were shared between the two low-altitude populations (25 expected; *p* = 0.0001), whereas in the high-altitude populations, 92 genes were shared out of 2240 (38 expected; *p* = 0.0001). (Fig 3B). We then conducted a Gene Ontology (GO) enrichment analysis to summarize the biological processes associated with these outlier genes. In contrast to individual loci, significantly enriched GO terms showed clear correlations among the low- and high-altitude populations (Fig 6). In the J1 and T1 populations, ‘leaf development’, ‘shoot system development’ and ‘response to bacterium’ were among the highest enriched GO categories, whereas the J3 and T4 outliers were enriched for terms ‘response to radiation’, ‘response to light stimulus’ and ‘cellular response to stress’.

**Figure 6.**
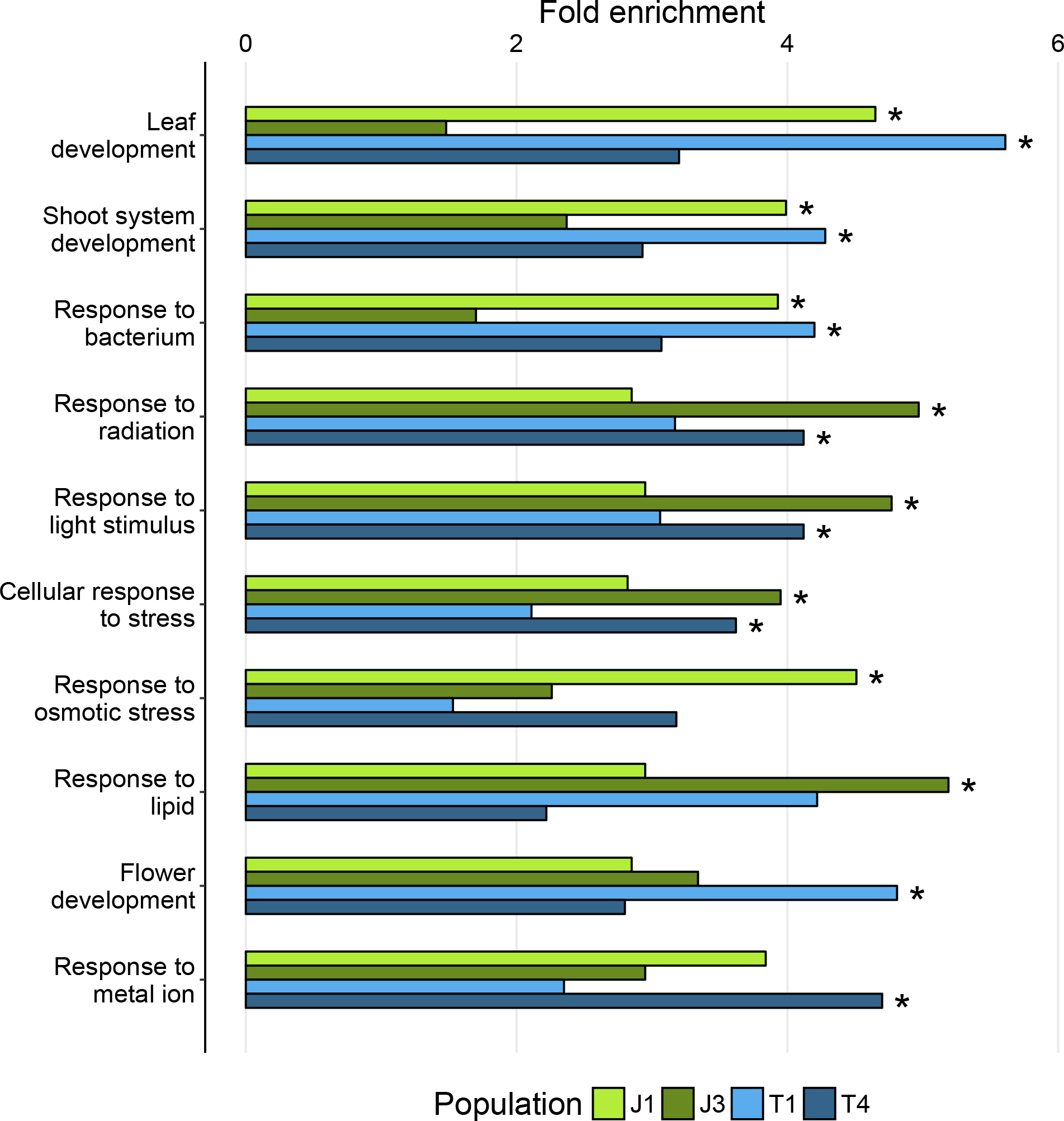
Highest enriched shared and unique GO terms with more than ten supporting genes among the significant (*q* < 0.05) PBS outliers. Terms with Bonferroni corrected *p*-value < 0.05 are marked with a star.

## Discussion

We found indications of lower genetic diversities and stronger population size contractions in the high-altitude populations than in the low-altitude populations. Combined with the low divergence time estimates from the demography models, these results point towards recent colonization of the alpine habitats in southwestern Norway. In contrast, populations in regions not influenced by the last glacial maximum (~20,000 years ago) likely reflect adaptation on a longer time scales [e.g. *A. halleri* in Japan (Kubota et al. 2015) and *A. thaliana* in Italy (Günther et al. 2016)], making our *A. lyrata* population set particularly suitable system for studying recent and reciprocal adaptation among populations connected by gene flow.

### Gene flow shapes patterns of neutral and adaptive variation

The effects of gene flow on genetic patterns of local adaptation have been intensely studied at the theoretical level (Lenormand and Otto 2000; Lenormand 2002; Griswold 2006; Kirkpatrick and Barton 2006; Bürger and Akerman 2011; Yeaman and Whitlock 2011; Akerman and Bürger 2014), but the empirical support for number of key predictions are still weak or missing. For example, under higher gene flow, is adaptation attributable to fewer large effect loci that are clustered in areas of lower recombination? We approach these questions by examining the lineage specific differentiation patterns among populations that receive variable number of migrants. The demography modelling indicated highly asymmetric gene flow between the neighbouring low- and high-altitude populations, which combined with the PBS analysis makes this setup more closely resemble a continent-island model (Bürger and Akerman 2011; Akerman and Bürger 2014) than a two-population model of local adaptation (Yeaman and Whitlock 2011; Akerman and Bürger 2014). Consequently, the low-altitude populations J1 and T1 showed PBS distributions with lower and less dispersed overall estimates than the high-altitude populations J3 and T4, and our simulation-based outlier detection model predicted more adaptive loci in the high-altitude populations. Our results further showed that outlier windows in the low-altitude populations were found in areas with lower recombination rates than in the high-altitude populations. The outliers were also slightly more densely clustered in the low-altitude populations, but the lack of correlation with recombination rates suggest that this pattern may be driven by other processes. In fact, if the genetic architecture underlying local adaptation evolves only through *de novo* mutations, a considerable amount of time may be needed for the signature patterns to form (Kirkpatrick and Barton 2006; Yeaman and Whitlock 2011; Yeaman 2013). As the observed differences have evolved recently, potentially during the last 1000 generations, patterns resistant to allele swamping (i.e. loci that in areas of low recombination and clustered loci) were likely present as standing genetic variation before the selection shift, leading to them being retained under gene flow independently. Additionally, these signals may in part be influenced by the interaction of gene flow and purifying selection, as alleles deleterious in one environment are more readily removed from areas of low recombination (Hudson and Kaplan 1995).

Similar results have previously been found among stickleback population pairs that exchange migrants at different rates (Marques et al. 2016; Samuk et al. 2017). However, by combining estimates of bidirectional gene flow with lineage specific selection analysis, we were able to examine these patterns not only between the Jotunheimen and Trollheimen study pairs, but also within them, making it possible to ascertain asymmetric effects and to better approach the genetic architecture underlying the local adaptation. Furthermore, although previous studies have shown a trend of outliers localizing in areas with lower than average recombination rates (Renaut et al. 2013), the effects of gene flow on this process have not (to our knowledge) been reported in any other plant taxa.

### Selection patterns suggest antagonistic pleiotropy between neighbouring populations

A major question in local adaptation research concerns the role of antagonistic pleiotropy in promoting adaptive divergence (Kawecki and Ebert 2004; Savolainen et al. 2013; Tiffin and Ross-Ibarra 2014; Wadgymar et al. 2017; Yoder and Tiffin 2017), and this issue is particularly relevant when the focus is on closely adjacent populations (Yeaman and Whitlock 2011; Akerman and Bürger 2014). A traditional way to search for genetic trade-offs is to measure presumably adaptive trait variation in contrasting environments and to find correlations between phenotypes and genotypes through genetic mapping (Fournier-Level et al. 2011; Ågren et al. 2013; Anderson et al. 2013; Leinonen et al. 2013). Although this approach has the potential advantage of linking phenotypes to fitness, focusing on preselected traits might cause important factors to be overlooked. Furthermore, the causative genes underlying the often-wide quantitative trait loci (QTL) intervals have rarely been discovered. Here, we used PBS scans to find loci showing patterns of opposing selection among the neighbouring low- and high-altitude populations, because under gene flow and unrestricted recombination, the frequency differentiation is only expected to be maintained by such fitness trade-offs.

Among the outliers potentially affected by antagonistic pleiotropy, we discovered one locus (*XRN2*) with clear evidence of allelic trade-off between the T1 and T4 populations. In *A. thaliana*, *XRN2* is known to be involved in various RNA processing tasks (Zakrzewska-Placzek et al. 2010), including posttranscriptional gene silencing (Gy et al. 2007); a defence response against viral mRNAs. Examination of nucleotide diversities and Tajima’s *D* estimates showed that the observed sequence patterns have likely resulted from two rapid and opposing selective sweeps, while the forward simulations indicated that selection has to be strong to produce these patterns in just under 600 generations (the upper confidence interval for the divergence time estimate). Furthermore, as suggested by our simulations, the highly negative Tajima’s *D* estimates are likely the result of a single or very few haplotypes being swept to a high frequency in each population. The sweeps might, however, have started from a standing genetic variation, but due to low *N*_*e*_ and high drift, nearly all haplotypes were lost during the initial selection phase, producing hard sweep like patterns (Orr and Betancourt 2001; Hermisson and Pennings 2017). In the absence of migration, a hard sweep with strong selection would likely lead to a wider footprint than observed here (Kaplan et al. 1989; Kim and Stephan 2002), but empirical studies have suggested that ongoing gene flow and selection against the migrant alleles might lead to a signal that is highly localised [e.g. selection on gene controlling colour patterns in deer mice (Linnen et al. 2013; Pfeifer et al. 2018)]. We nevertheless acknowledge that this signal could more easily arise from selection that is older than our demography model suggests, but given the recent postglacial colonization of Scandinavia by *A. lyrata* (<10,000 generations ago) (Mattila et al. 2017), a genetic trade-off accompanied by gene flow is still the most likely source for the observed patterns. Reduced recombination, on the other hand, is likely not a factor here, because our linkage map and the short area influenced by the selective sweeps indicate that this area of the chromosome recombines freely.

As shown by Martin & Lenormand (2006, 2015), conditional neutrality may actually be the dominant driver of local adaptation across all environments, because it can evolve through purely deleterious mutations (as opposed to antagonistic pleiotropy, which requires a beneficial effect in at least one environment). This fact is probably reflected in our results as well, as the footprint of background selection can sometimes be similar to genetic hitchhiking (Comeron 2017), but some trade-off loci are likely also identified as conditionally neutral due to lack of power to detect selection in both lineages (Anderson et al. 2013; Tiffin and Ross-Ibarra 2014; Wadgymar et al. 2017; Yoder and Tiffin 2017). The divergence between these populations might also be too recent for antagonistic pleiotropy to become more common, as it is a prediction based on longer evolutionary time scales (Yeaman and Whitlock 2011). Studies based on QTL mapping have suggested that genetic trade-offs may be more abundant between the long since diverged Swedish and Italian *A. thaliana* populations (Ågren et al. 2013; Price et al. 2018). However, in the absence of gene flow, these patterns are not due to interaction of migration and selection, but they might e.g. reflect genomic constraints. In any case, the apparent prevalence of conditional neutrality can result in freer spread of alleles, which might be beneficial for these *A. lyrata* populations upon climate change (Fournier-Level et al. 2011; Hämälä et al. 2018).

### Resistance to solar radiation is selected for in alpine environments

Besides genome-wide patterns of adaptive differentiation, our results revealed genes and biological process that may underlie colonization of alpine and lowland habitats in southwestern Norway. A major factor associated with highland environments is the increase in solar radiation intensity (Körner 2007). Indeed, earlier studies conducted on plants (Fischer et al. 2013), as well as on humans (Yang et al. 2017), have detected directional selection on UV resistance genes in high-altitude populations. Our analysis also discovered a UV resistance gene *TT5* (Li et al. 1993) among the most significant outliers in J3 and a DNA break repair (a trait related to increased radiation levels) gene *RAD23C* (Farmer et al. 2010) was found in T4. Furthermore, genes involved in response to light stimulus were discovered in both populations, and the highest shared outlier between J3 and T4 was *ADG1*; a gene thought to be important in acclimation to high light intensities in *A. thaliana* (Schmitz et al. 2014). As indicated by our GO enrichment analysis, loci involved in ‘response to radiation’ were more numerous than expected among the outlier loci in the two alpine populations. Additional highly enriched GO categories were ‘response to light stimulus’ and ‘cellular response to stress’, which might also be linked to adaptation under increased solar radiation. Selection on radiation resistance genes may not, however, be ubiquitous among highland populations, as shown by studies on *A. halleri* in Japan (Kubota et al. 2015) and on *A. thaliana* in Italy (Günther et al. 2016). Therefore, these results suggest that solar radiation imposed selection can be a significant driver of adaptive divergence in the northern latitudes, even though the high-altitude areas in Scandinavia are relatively low elevation compared to highest regions in the world.

### Vegetative growth and bacterial defence are important traits in lowland adaptation

We also discovered interesting selection patterns in the low-altitude populations. Genes involved in leaf development and vegetative growth were found among the top outliers in J1 and T1, and the corresponding ‘leaf development’ GO category was significantly enriched in both gene sets. ‘Shoot system development’, another significantly enriched term, is especially interesting because we have previously shown that lowland populations generally produce longer flowering shoots than alpine populations, and the trait had a positive correlation with fitness at the Finnish sea level field site (Hämälä et al. 2018). Bacterial defence seems to be another important trait in the lowland habitats. The GO term ‘response to bacterium’ was significantly enriched in both lowland populations, and the top outlier in J1 (*SRT2*) is involved in that process (Wang et al. 2010). Selection on bacterial defence genes has previously been discovered among populations from different altitudes in *A. halleri* (Fischer et al. 2013; Kubota et al. 2015) and in *A. thaliana* (Günther et al. 2016), but as those studies relied on traditional divergence outlier methods, the authors could not infer whether the selection target had been the low- or the high-altitude lineage. Furthermore, *XRN2*, the gene exhibiting allelic trade-off between T1 and T4, is thought to be involved in immune responses in *A. thaliana* (Gy et al. 2007), but it also has a role in general RNA processing (Zakrzewska-Placzek et al. 2010), so the causal factor behind the trade-off may not be related to pathogen defence.

Despite the fact that many genes with same biological processes were shared between the two high- and low-altitude populations, almost all individual loci were population specific. This lack of correlation at the gene level likely indicates that populations living in similar environments have responded to same environmental pressures through different genetic pathways, suggesting either subtle differences in phenotypes under selection or that selection has acted on different genes underlying the same polygenic traits. Considering the recent divergence time estimate between populations from the two alpine areas (~2800 years ago) and the growing evidence of convergent local adaptation even among more distantly related groups (Arnold et al. 2016; Yeaman et al. 2016), these results suggest that adaptation to altitude specific environments in *A. lyrata* is not constrained to only few key genes, which might further aid adaptation to future climates.

## Conclusions

By studying recently diverged, phenotypically differentiated and locally adapted *A. lyrata* populations, we have gained novel insights into adaptive processes driving differentiation under ongoing gene flow. As predicted by theory, selection outliers in the lowland populations were located in areas with lower recombination rates than in the alpine populations. We also found a locus displaying clear footprints of strong opposing selective sweeps in the Trollheimen populations; a pattern likely caused by antagonistic pleiotropy. However, most selected loci showed indications of conditional neutrality, potentially reflecting the recent divergence between these populations. Phenotypes associated with the outliers further revealed biological processes that may underlie recent alpine and lowland adaptation in northern Europe. These results contribute to understanding of processes driving adaptive differentiation under gene flow, as well as of traits and biological processes underlying alpine adaptation in northern latitudes.

## Materials and methods

### Study populations

We studied altitude adaptation among Norwegian *Arabidopsis lyrata* populations from two alpine areas (Jotunheimen and Trollheimen). Both areas were represented by one low- and one high-altitude population (Fig 1; Table S1). Based on patterns of microsatellite variation, populations within both alpine areas form distinct genetic clusters (Gaudeul et al. 2007). The fitness and phenotypic variation of these populations were previously studied in Hämälä et al. (2018). In that study, the areas were represented by four populations, which were abbreviated as J1 to J4 and T1 to T4. Here, we retain the naming convention and call the populations J1 (300 m.a.s.l), J3 (1,100 m.a.s.l), T1 (10 m.a.s.l) and T4 (1360 m.a.s.l). The distance between the J1 and J3 growing sites is approximately 25 km, whereas the approximate distance between T1 and T4 is 30 km. The two alpine areas are roughly 100 km apart (Fig 1). We previously showed that individuals from J1 and J3 exhibited local superiority when grown at reciprocal common garden sites in Norway. The T1 and T4 individuals were not planted at their local environments, but when grown at common garden in Finland, they expressed phenotypic differences consistent with altitude adaptation (Hämälä et al. 2018).

### Whole-genome sequencing

Whole-genome data from two previously published studies were used (Mattila et al. 2017; Hämälä et al. 2018): nine individuals from J1, 12 from J3 and five from T1. For the present study, we also sequenced nine individuals from T4 population, which exhibited more high-altitude specific phenotypes (earlier flowering start and shorter flowering shoots, as well as higher fruit production) in our field experiments than the previously sequenced T3 population (Hämälä et al. 2018). In addition, a German population (abbreviated as GER) consisting of six individuals was used as an outgroup in the selection scan (see ‘Selection scan’) and the German and a Swedish population (SWE; consisting also of six individuals) were used as comparison groups in admixture and principal component analysis (see ‘Analysis of genetic diversity and population structure’). The German and Swedish individuals, as well as five individuals from J3, came from Mattila et al. (2017), whereas the other previously sequenced individuals were from Hämälä et al. (2018). In all three studies, DNA was extracted from fresh leaves using NucleoSpin Plant II kits (Macherey-Nagel), the libraries for Illumina whole-genome sequencing were prepared with NEBNext master mix kits (New England Biolabs), and the sequencing was done with Illumina HiSeq2000 (Mattila et al. 2017) and HiSeq2500 (Hämälä et al. 2018; this study) in Institute of Molecular Medicine Finland, University of Helsinki, using PE100 chemistry. The median read coverage per individual ranged from six to 25. The low coverage in some individuals combined with the variable nature of the short-read sequencing means that information contained in large part of the data set is insufficient to call genotypes with high confidence (Nielsen et al. 2011). Therefore, to lessen the bias caused by uncertain genotypes, we adopted a SNP call free approach and based all estimated statistics on genotype likelihoods.

### Sequence processing and allele frequency estimation

Low quality reads and sequencing adapters were removed using Trimmomatic (Bolger et al. 2014). The reads were aligned to *A. lyrata* v1.0 reference genome (Hu et al. 2011) with BWA-MEM (Li and Durbin 2009). Duplicated reads were removed and indels realigned with GATK (DePristo et al. 2011). Likelihoods for the three possible genotypes in each biallelic site were then calculated with the GATK model in ANGSD (Korneliussen et al. 2014). We only used reads with mapping quality over 30, while sites needed to have quality over 20 and sequencing depth no less than 4×. Allele frequencies for the selection scan were then estimated directly from the genotype likelihoods using a maximum likelihood model by Kim et al. (2011). The strict filtering associated with SNP calling (commonly the ranking genotype needs to be ten times more likely than the others) can especially reduce the number of heterozygote calls in areas of low coverage (for a comparison between SNP calls and genotype likelihoods in our data, see Fig S12). The method used here circumvents that problem by taking the genotype uncertainty into account, producing unbiased allele frequency estimates even with the minimum threshold coverage (Kim et al. 2011; Nielsen et al. 2011).

### Analysis of genetic diversity and population structure

We studied genetic variation within populations by estimating two summary statistics with ANGSD (Korneliussen et al. 2014): nucleotide diversity *π*, which is a measure of the population mutation rate *θ* = 4*N*_*e*_*μ* (Nei and Li 1979), and Tajima’s *D*, which approximates how far the population is from a mutation-drift balance (Tajima 1989). Principal component analysis (PCA) and admixture analysis were conducted to further assess the genetic relationships between the study populations. Genotype likelihoods were estimated for 4-fold degenerate sites with ANGSD and used as a input for PCAngsd (Meisner and Albrechtsen 2018). As suggested by the program documentation, we used the optimal number of PCs to define the likely number of ancestral populations (K).

### Demography simulations

Site frequency spectra based coalescent simulations were used to estimate divergence times, migration rates and effective population sizes. Estimates involving the Jotunheimen populations J1 and J3 were inferred as part of an earlier study (Hämälä et al. 2018), and here we conducted additional simulations for the Trollheimen populations T1 and T4. Briefly, derived site frequency spectra (SFS) were estimated for 4-fold degenerate sites in ANGSD (Korneliussen et al. 2014) and the demography models were fitted to these in fastsimcoal2 (Excoffier et al. 2013). We tested four different migration models between the Norwegian populations: no migration, unidirectional migration (from 1 to 2 and from 2 to 1) and bidirectional migration. To explore alternative explanations for the estimated migration parameters, we also tested whether asymmetric expansion after a bottleneck may lead to spurious signals that could be mistaken for gene flow. Simulation were repeated 50 times to acquire global maximum likelihood estimates for the parameters. Model selection was based on the Akaike information criterion (AIC) scores. We then used 100 nonparametric bootstrap SFS to define 95% confidence intervals for the parameter estimates. For more information about the demography analysis, see Hämälä et al. (2018).

As our approach for examining the genetic architecture of local adaptation rely on accurately estimating the direction of gene flow, we evaluated the performance of the inference method with simulated data. We used MLEs from the best supported demography models (Table S4) to generate 10,000 × 10 Kb fragments for each population. As with the observed data, SFS from the simulated samples were estimated with ANGSD and used as an input for fastsimcoal2. A reasonable correspondence between parameters estimates from the observed and simulated data sets (Table S5) confirmed that fastsimcoal2 performs reliably with our *A. lyrata* populations.

### Selection scan

Selected sites were detected by scanning the chromosomes for areas of unexpectedly high differentiation between the populations; a pattern indicative of directional selection (Lewontin and Krakauer 1973). The relative levels of differentiation were estimated with *F*_ST_ measure by Hudson et al. (1992): 1 – (*H*_W_ / *H*_B_), where *H*_W_ and *H*_B_ are the mean number of differences within and between populations, respectively. However, here we used a more specific formula, developed for the two population, two allele case by Bhatia et al. (2013):

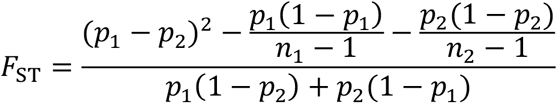

where *n*_*i*_. is the sample size and *p*_*i*_. is the minor allele frequency in the two populations to be compared. This estimator has been shown to be unbiased by unequal sample sizes and less prone to overestimating differentiation than other commonly used *F*_ST_ measures (Bhatia et al. 2013). The selection scan was conducted in SNP-based window sizes to prevent biasing the estimates with unequal SNP numbers. As the average linkage disequilibrium decays rapidly in these *A. lyrata* populations (Fig S13), 50-SNP non-overlapping sliding windows (median length ~3 Kb) provided a good trade-off between resolution and independence of the estimates. We also used 1-SNP and 150-SNP (median length ~10 Kb) windows to confirm patterns related to genetic architecture of local adaptation. *F*_ST_ for a window of size *n* was calculated using the weighting method by Reynolds et al. (1983): 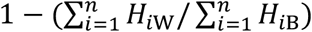. For lower number of markers (such as within windows), this approach is often more reliable than averaging the *F*_ST_ estimates over loci (Weir and Hill 2002; Bhatia et al. 2013). *F*_ST_ can, however, be inflated by reduced within-population nucleotide diversities, brought on e.g. by background selection or lowered recombination (Charlesworth 1998; Cruickshank and Hahn 2014). For this reason, we also estimated absolute levels of differentiation between the populations; an index commonly called *d*_XY_ (Nei 1987; Cruickshank and Hahn 2014). To make it compatible with allele frequencies estimated from genotype likelihoods, *d*_XY_ for a window of size *n* was calculated by simply excluding the within-populations component of *F*_ST_:

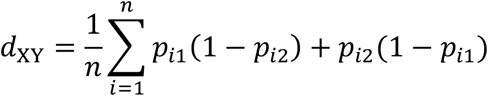

This measure is indifferent to within-population levels of diversity, but it can be biased by unequal sample sizes. Therefore, to balance the shortcomings of both *F*_ST_ and *d*_XY_, we only considered sites that were detected as outliers using both measures.

A single differentiation measure, either *F*_ST_ or *d*_XY_, can detect localized selection, but it cannot distinguish which lineage has been the target. A recently developed method, population branch statistic (PBS) (Yi et al. 2010), overcomes this limitation by comparing differentiation estimates between two closely related populations and an outgroup. The *F*_ST_ and *d*_XY_ values were first transformed into relative divergence times: *T* = −ln(1 − *X*), where *X* is the differentiation measure. PBS for population 1 was then estimated as:

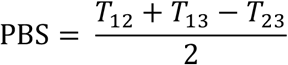

The obtained value quantifies the magnitude of allele frequency change in a lineage 1 since its divergence from the closely related population 2 and the outgroup 3. Selection acting only on a single lineage would appear as higher than neutral differentiation between the focal populations (i.e. *T*_12_) and one of the outgroup comparisons (e.g. *T*_13_), whereas different alleles being under selection in both focal populations would lead to high differentiation between all three population comparisons (i.e. *T*_12_, *T*_13_ and *T*_23_). By choosing a sufficiently different outgroup, we can reduce the chance that loci under differential selection between the focal populations are also under selection in the outgroup, as this can introduce bias into the analysis. Therefore, the German population was chosen as the outgroup over the Swedish one, because it diverged from the Norwegian populations around 30,000 generation ago (Table S4) and its natural growing environment is highly different from the Norwegian environments (Leinonen et al. 2009). Lineage specific selection patterns among the focal populations were estimated by calculating PBS estimates for population trios J1-J3-GER and T1-T4-GER. To further evaluate if selection acting on the outgroup lineage influences our results, we compared population specific PBS outliers (for explanation of the outlier detection, see next section) to *F*_ST_ and *d*_XY_ outliers estimated between the corresponding focal population and the outgroup (e.g. PBS outliers in population J1 were compared against regular differentiation outliers between J1 and GER). Each comparison had minimal (~5%) overlap, suggesting that selection acting on the outgroup lineage causes no major bias in categorization of the putatively adaptive loci.

### Outlier detection

We compared the PBS estimates against simulated samples to find sites that show higher differentiation than expected under neutrality. Neutral data were generated with coalescent models in ms (Hudson 2002), by taking into account the genome-wide recombination rates and the demographic history of these populations. This approach can produce realistic approximations of the null distribution, which generally leads to fewer false positives compared to methods based on specific population genetic or statistical models (Lotterhos and Whitlock 2015; Hoban et al. 2016). The relevant MLEs from the demography models (i.e. divergence times, migration rates and effective population sizes, going back to a most recent common ancestor of these populations) were used for each population comparison. Recombination rates for sequences that corresponded in size to observed window lengths were pulled randomly from a linkage map published in an earlier study (Hämälä et al. 2017). The map was constructed by crossing populations representing the two *A. lyrata* subspecies (one from Eurasia and one from North America) and it is not specific for the populations of the present study. However, we expect that in *A. lyrata*, as in many other species (Ritz et al. 2017), the large-scale recombination patterns are conserved among populations, making the linkage map a suitable tool for the current study. The mutation rate was set to 7×10^−9^ following Ossowski et al. (2010). Using the simulated data, we acquired ~50,000 neutral PBS estimates for each population, which constituted the null distributions for outlier testing. We defined *p*-values as the proportion of neutral estimates that had the same or higher PBS value than the observed one. The *p*-values were subsequently transformed into false discovery rate based *q*-values (Storey and Tibshirani 2003) to reduce the bias caused by multiple testing. PBS estimates with *q*-value lower than 0.05 were considered significant. We have implemented this PBS scan and outlier detection method into a new C program, PBScan, available at: https://github.com/thamala/PBScan

### Analysis of putatively adaptive loci

To examine processes potentially affecting localization of the adaptive loci, we estimated recombination rates, gene densities, and mutation rates for the outlier areas. We used the linkage map and *A. lyrata* v1.0 genome annotation (Hu et al. 2011) to estimate recombination rates and gene densities, respectively. Additionally, we used *A. lyrata* – *A. thaliana* whole-genome alignments from Mattila et al. (2017) to estimate the proportion of synonymous nucleotide substitutions per synonymous site (*d*_S_), which served as a proxy for mutation rate (Begun and Aquadro 1992). To evaluate whether outlier statistics differ between populations, distributions were compared with bootstrap based Kolmogorov-Smirnov test (Sekhon 2011) with 10,000 replicates.

For one especially interesting locus, *XRN2*, we also examined what evolutionary scenarios may have produced the selection patterns by conducting simulations under different forward genetic models in SLiM 2 (Haller and Messer 2017). As in the case of neutral simulations, we used the estimated demography parameters, mutation rate from Ossowski et al. (2010), and recombination information from the corresponding area of the linkage map (*r* = 3.7×10^−8^). Selection parameters (4*N*_*e*_*s* = 100, 1000 and 10,000) were based on Pennings and Hermisson (2006). Code for the SLiM 2 simulations is available at: https://github.com/thamala/SLiM2_scripts

Lastly, we examined to what biological processes these outliers are associated with using a Gene Ontology (GO) (Ashburner et al. 2000) analysis. Although GO enrichment analysis has known caveats, such as those related to difficulty in defining the null expectations for significant testing or spurious signals caused by linked selection, it remains one of the best tools for summarizing functions among large quantities of genes (Gaudet and Dessimoz 2017). PANTHER tools (Mi et al. 2017) was used to detect significantly enriched terms among genes that localized within or were closer than 5 Kb of significant (*q*-value < 0.05) outlier windows.

## Acknowledgements

We thank H. Stenøien for sharing information about the *A. lyrata* growing sites in Norway, T. Toivainen and J. Tyrmi for sample collections, and T. Mattila for providing part of the data set. We are also grateful for P. Ingvarsson and members of the Oulu Plant Genetics Group for valuable discussions and comments on the manuscript. IT Center for Science (CSC) supplied computational resources. This work was supported by Biocenter Oulu (to T.H and O.S), Eemil Aaltonen Foundation (to T.H) and Academy of Finland’s Research Council for Biosciences and Environment (decision 132611 to O.S).

## Supplementary Information

**Figure S2.**
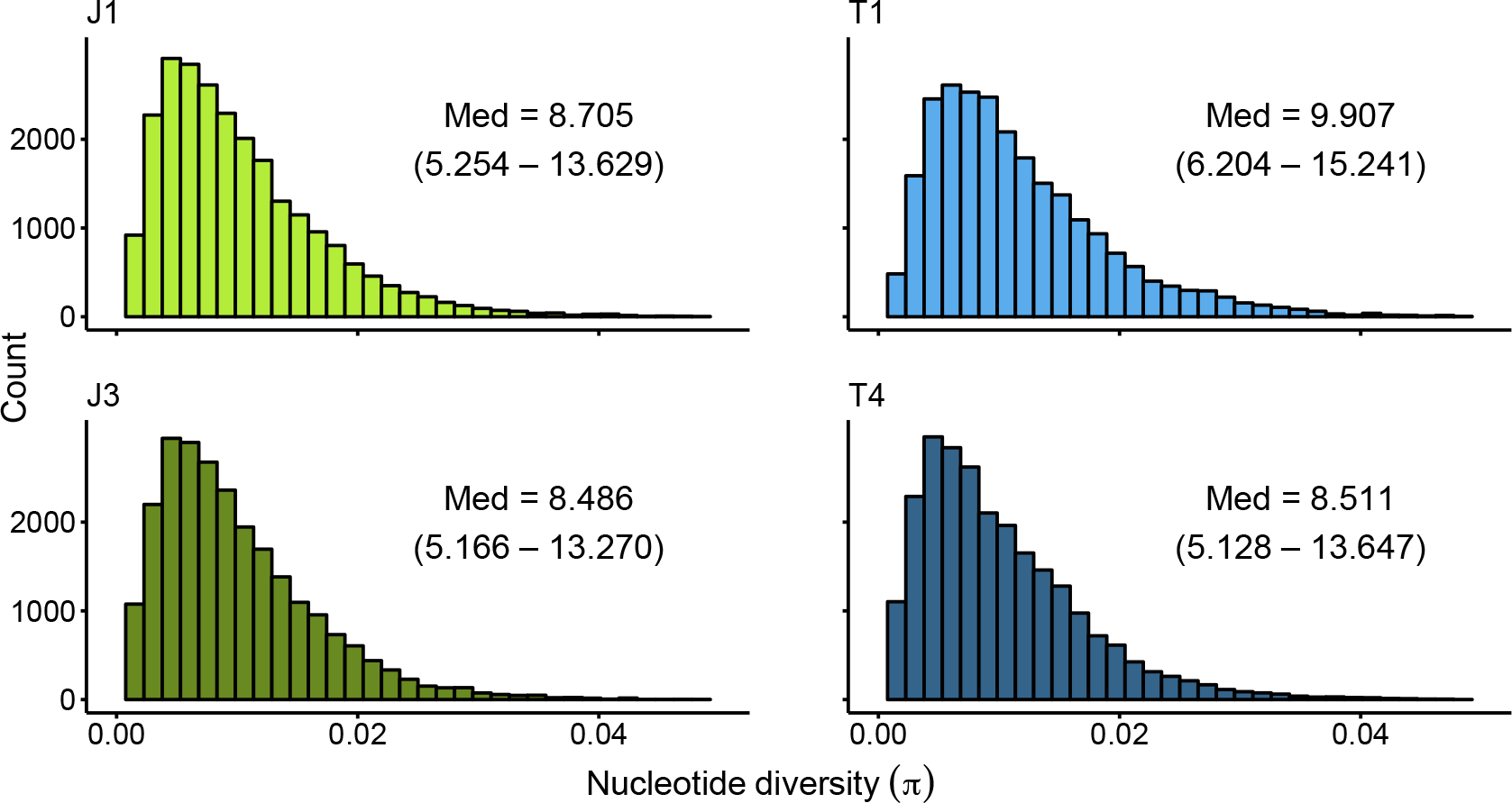
Whole-genome distributions of nucleotide diversity (*π*). Median estimates and interquartile ranges (in the units of ×10^−3^) are marked in each figure. Estimated in 50 Kb non-overlapping sliding windows. The low-altitude populations have *π* estimates that are statistically higher than in the high-altitude populations (J1-J3 *p*-value = < 2.2×10^−16^, T1-T4 *p*-value < 2.2×10^−16^; one-sided bootstrap Kolmogorov-Smirnov [KS] test).

**Figure S3.**
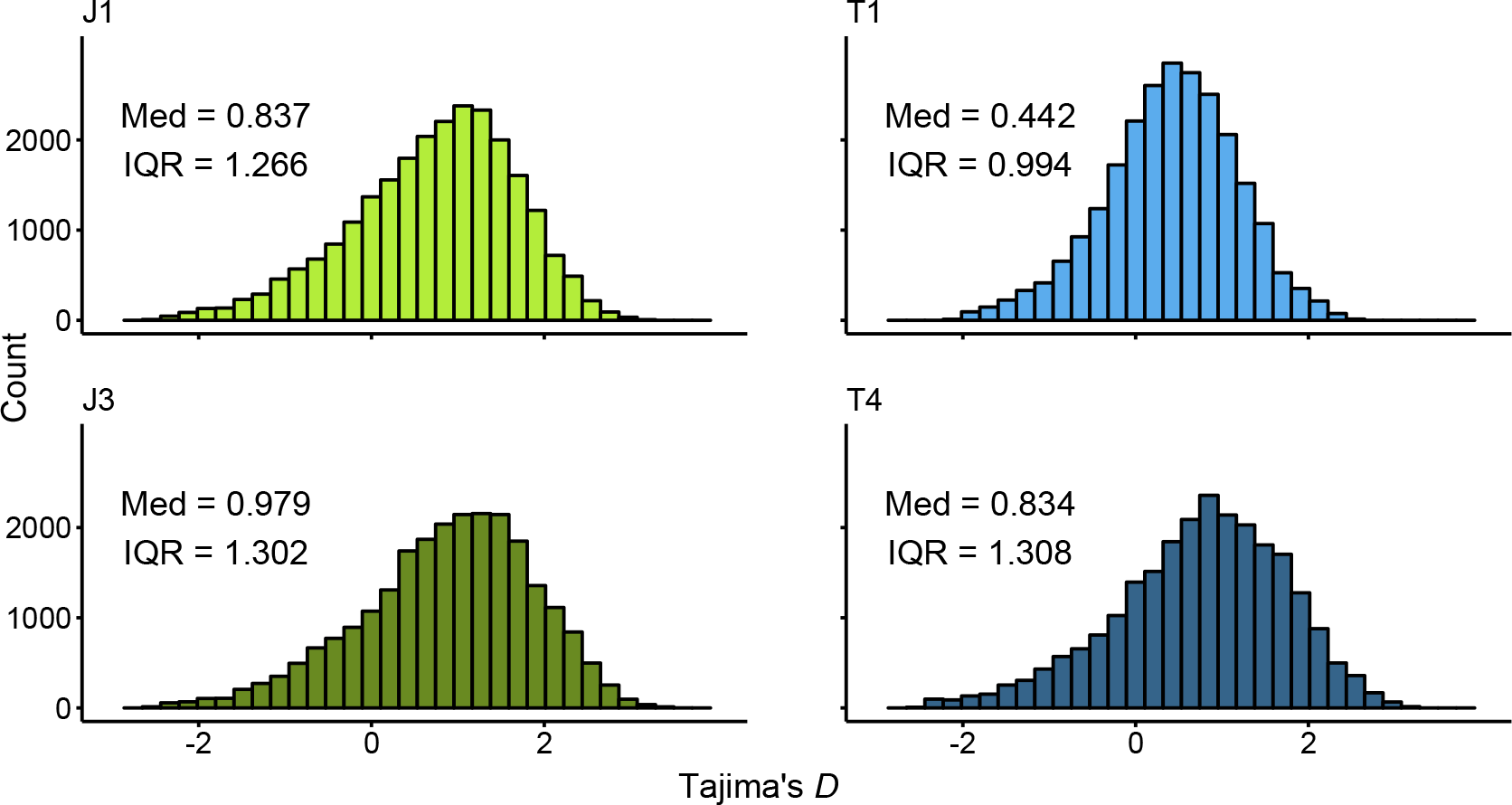
Whole-genome distributions of Tajima’s *D*. Median estimates and interquartile ranges are marked in each figure. Estimated in 50 Kb non-overlapping sliding windows. The low-altitude populations have *D* estimates that are statistically lower than in the high-altitude populations (J1-J3 *p*-value < 2.2×10^−16^, T1-T4 *p*-value < 2.2×10^−16^; one-sided bootstrap KS test).

**Figure S4.**
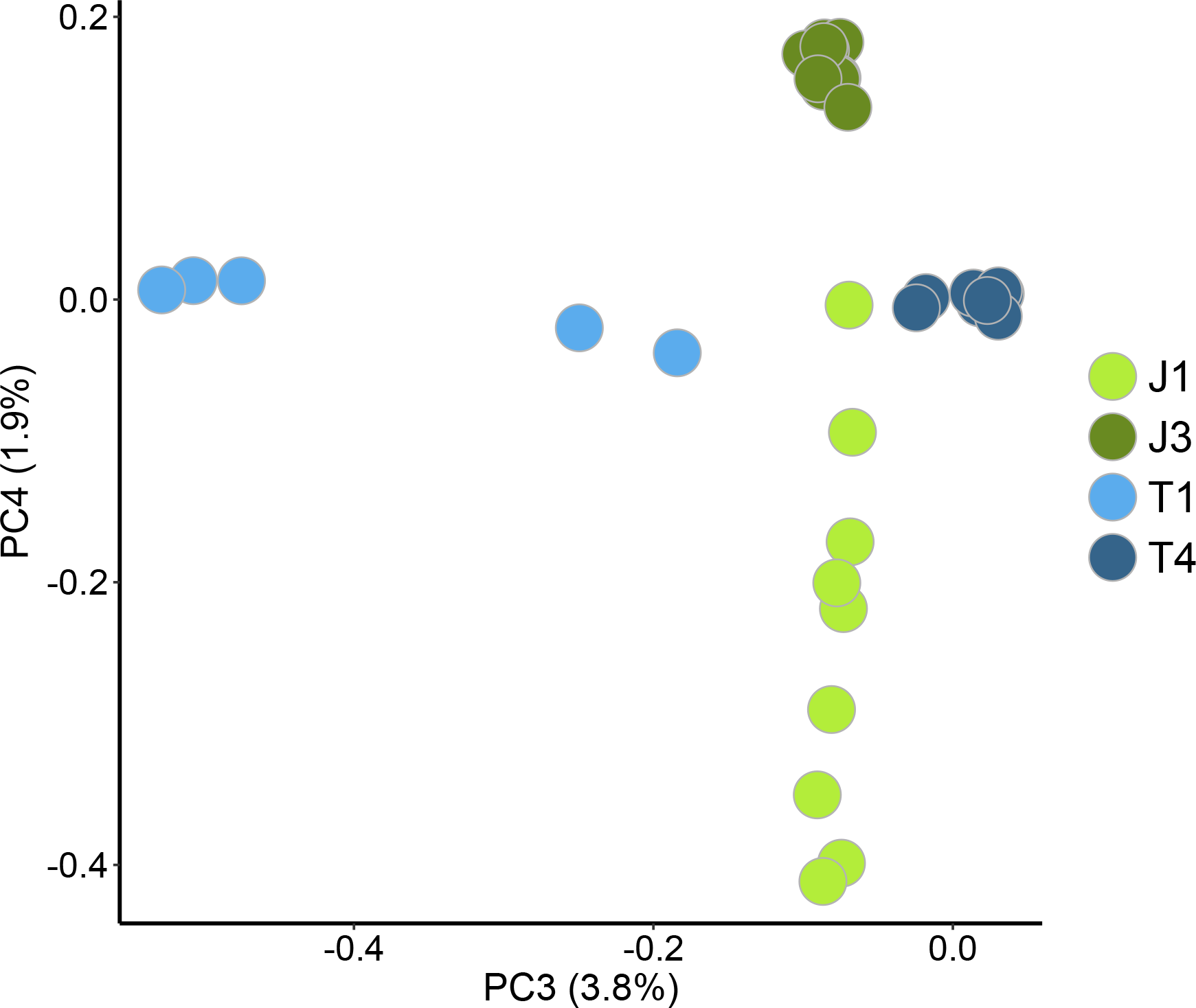
Variation along the third and fourth principal components in the Norwegian-only PCA.

**Figure S4.**
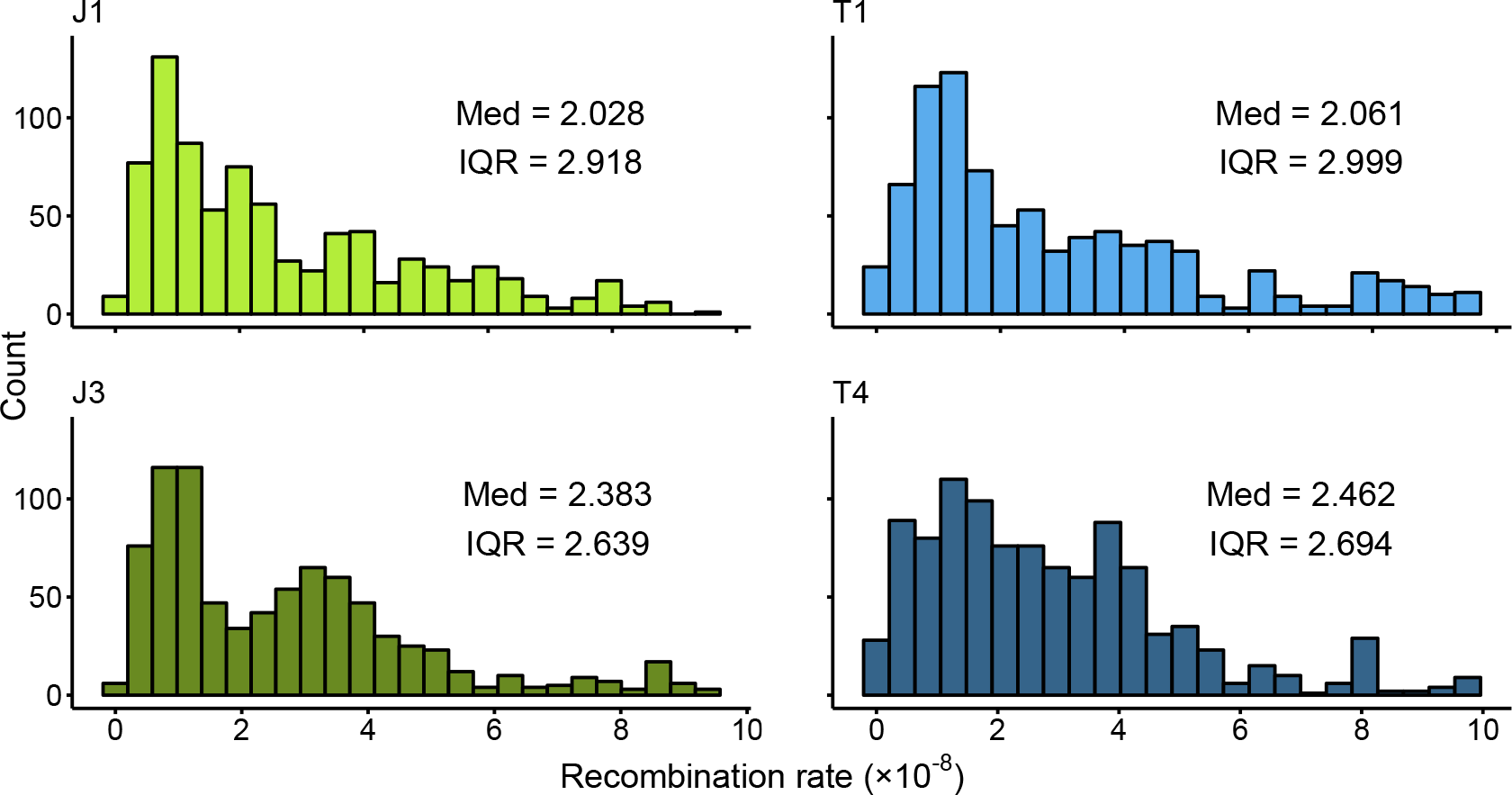
Per base pair recombination rate distributions for the PBS outlier areas. Median estimates and interquartile ranges are marked in each figure. The low-altitude populations have recombination rate estimates that are statistically lower than in the high-altitude populations (J1-J3 *p*-value = 0.005, T1-T4 *p*-value = 0.0001; one-sided bootstrap KS test).

**Figure S5.**
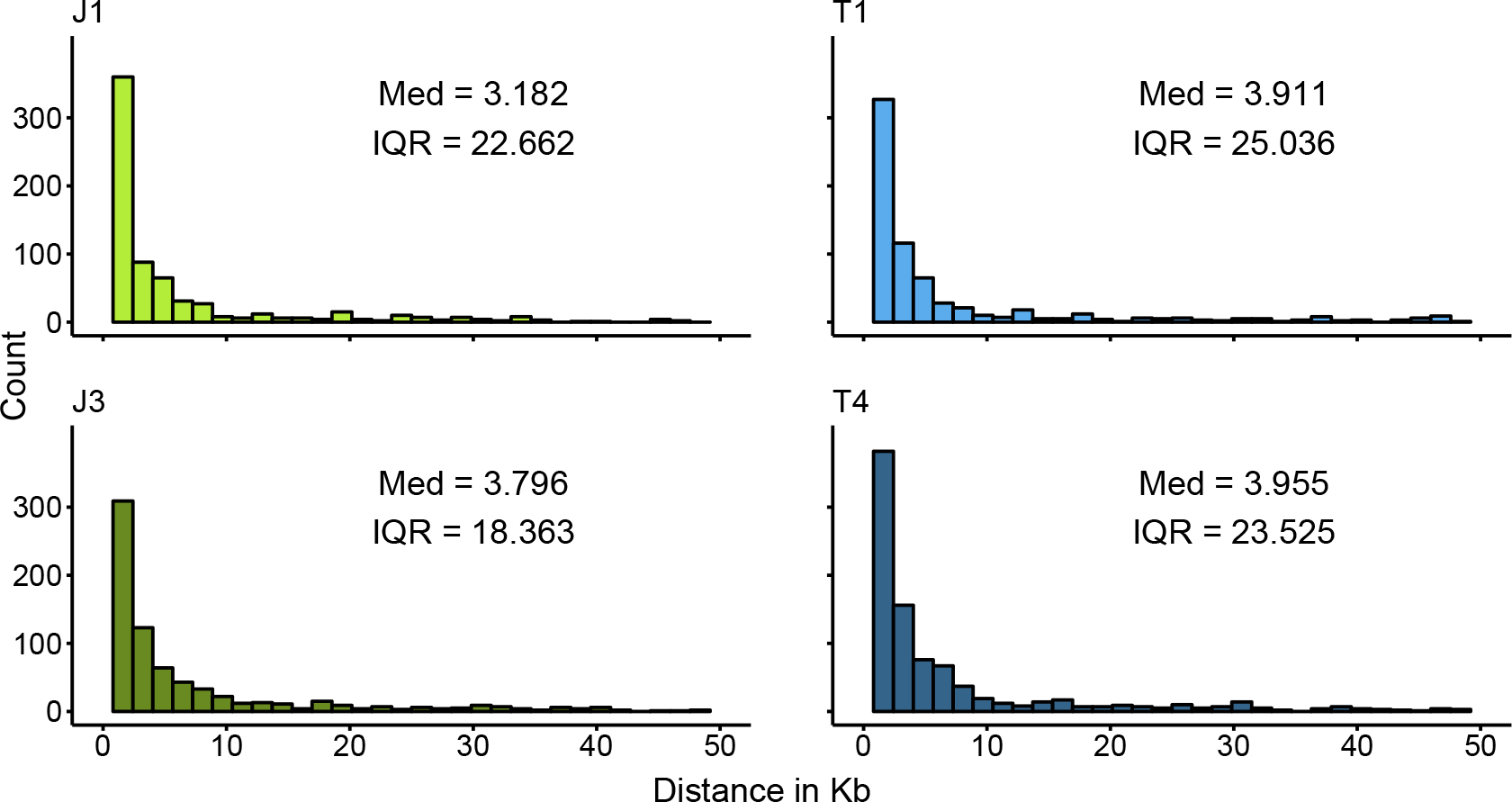
Distributions for between PBS outlier windows. Median estimates and interquartile ranges are marked in each figure. The low-altitude populations have distance estimates that are statistically lower than in the high-altitude populations (J1-J3 *p*-value = 0.01, T1-T4 *p*-value = 0.006; one-sided bootstrap KS test).

**Figure S6.**
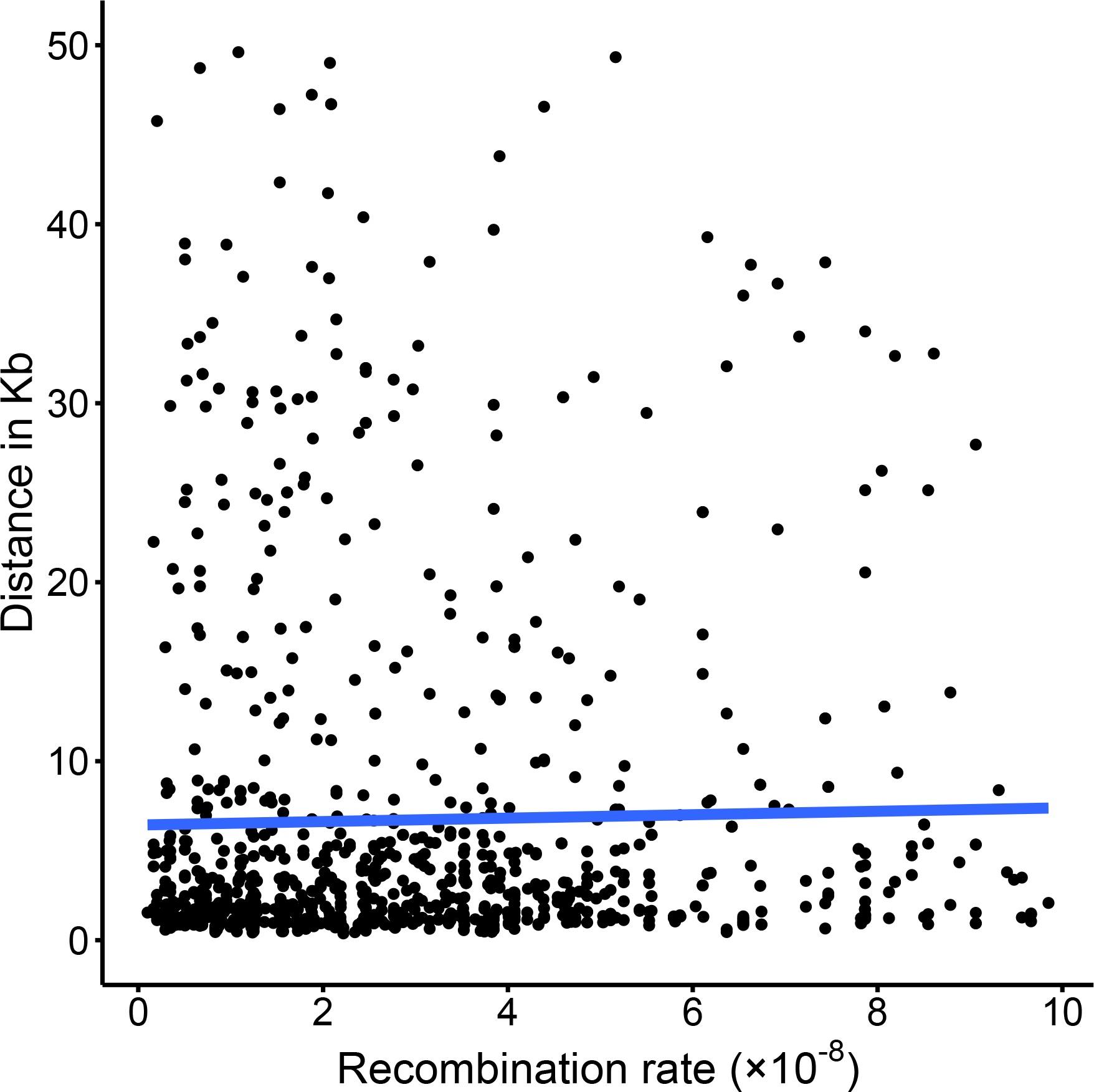
Correlation among within outlier window recombination rates and between outlier window distances. Pearson *r* = 0.022, *p* = 0.482.

**Figure S7.**
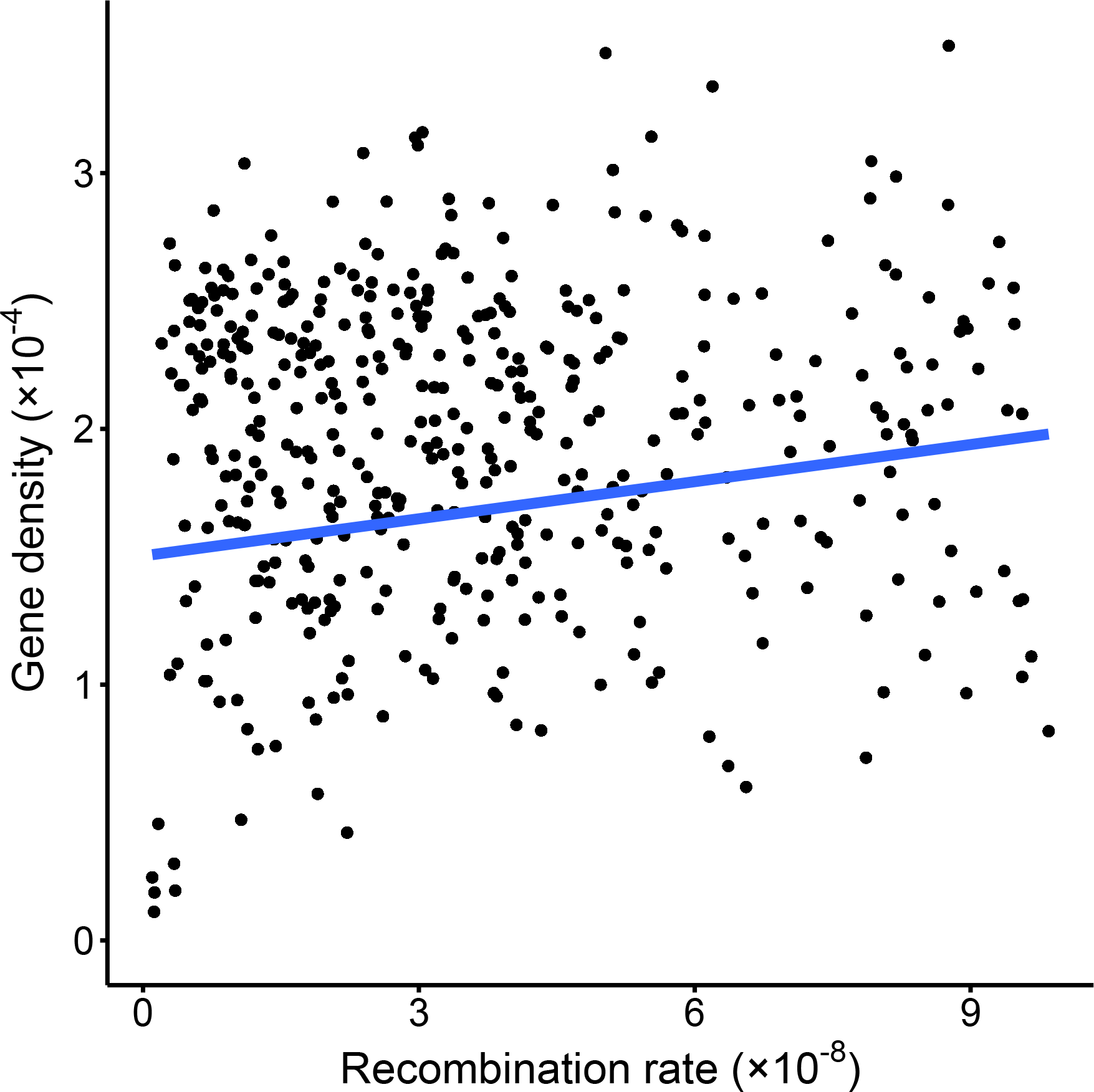
Genome-wide correlation among per base pair recombination rates and per base pair gene densities. Pearson *r* = 0.159, *p* < 2.2×10^−16^.

**Figure S8.**
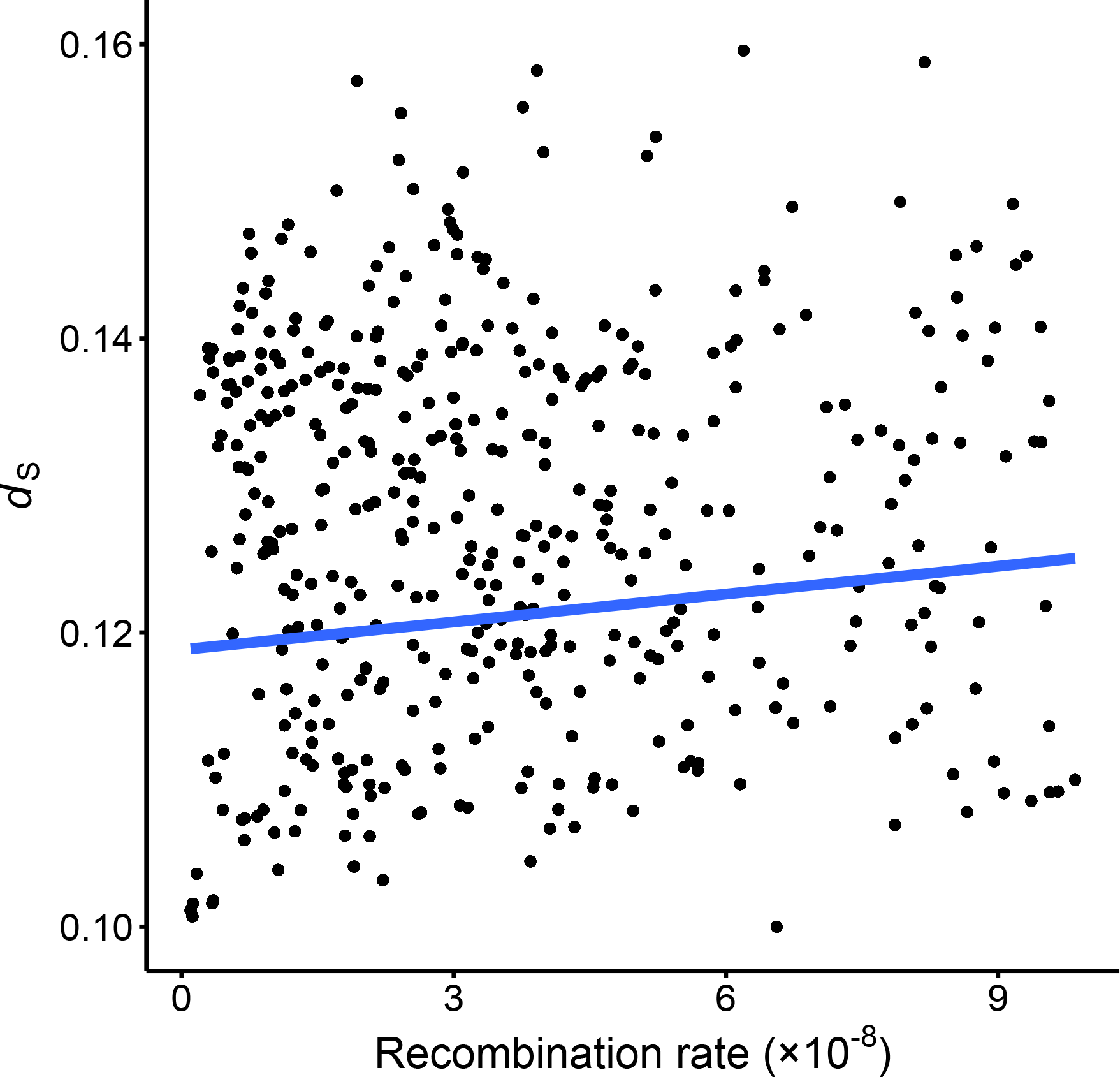
Genome-wide correlation among per base pair recombination rates and the rate of synonymous nucleotide substitutions (*d*_S_). Pearson *r* = 0.109, *p* < 2.2×10^−16^.

**Figure S9.**
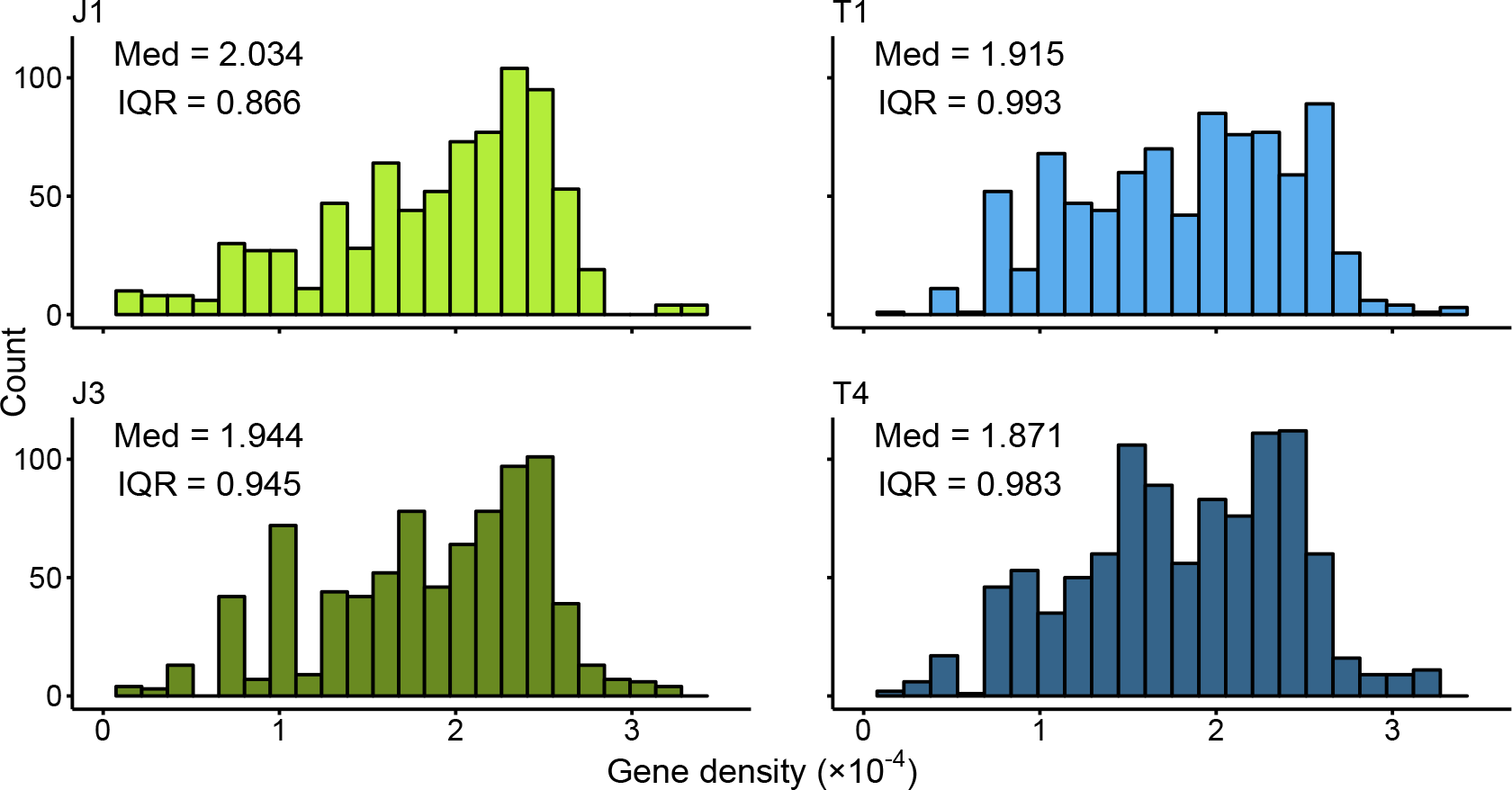
Per base pair gene density distributions for the PBS outlier areas. Median estimates and interquartile ranges are marked in each figure. The gene density estimates do not differ statistically between the low- and high-altitude populations (J1-J3 *p*-value = 0.08, T1-T4 *p*-value = 0.16; two-sided bootstrap KS test).

**Figure S10.**
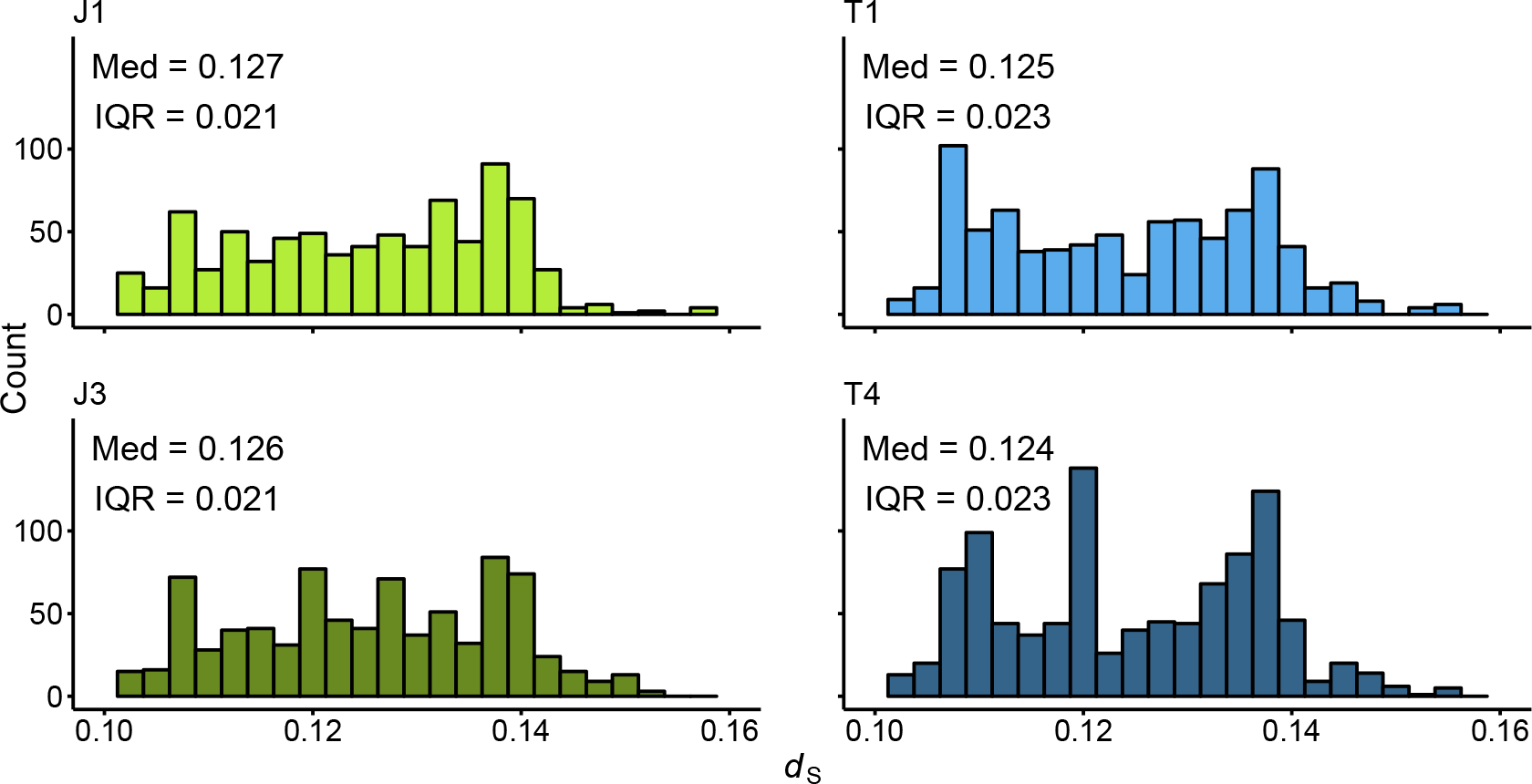
Synonymous nucleotide divergence distributions (*d*_S_) for the PBS outlier areas. Median estimates and interquartile ranges are marked in each figure. The *d*_S_ estimates do not differ statistically between the low- and high-altitude populations (J1-J3 *p*-value = 0.245, T1-T4 *p*-value = 0.212; two-sided bootstrap KS test).

**Figure S11.**
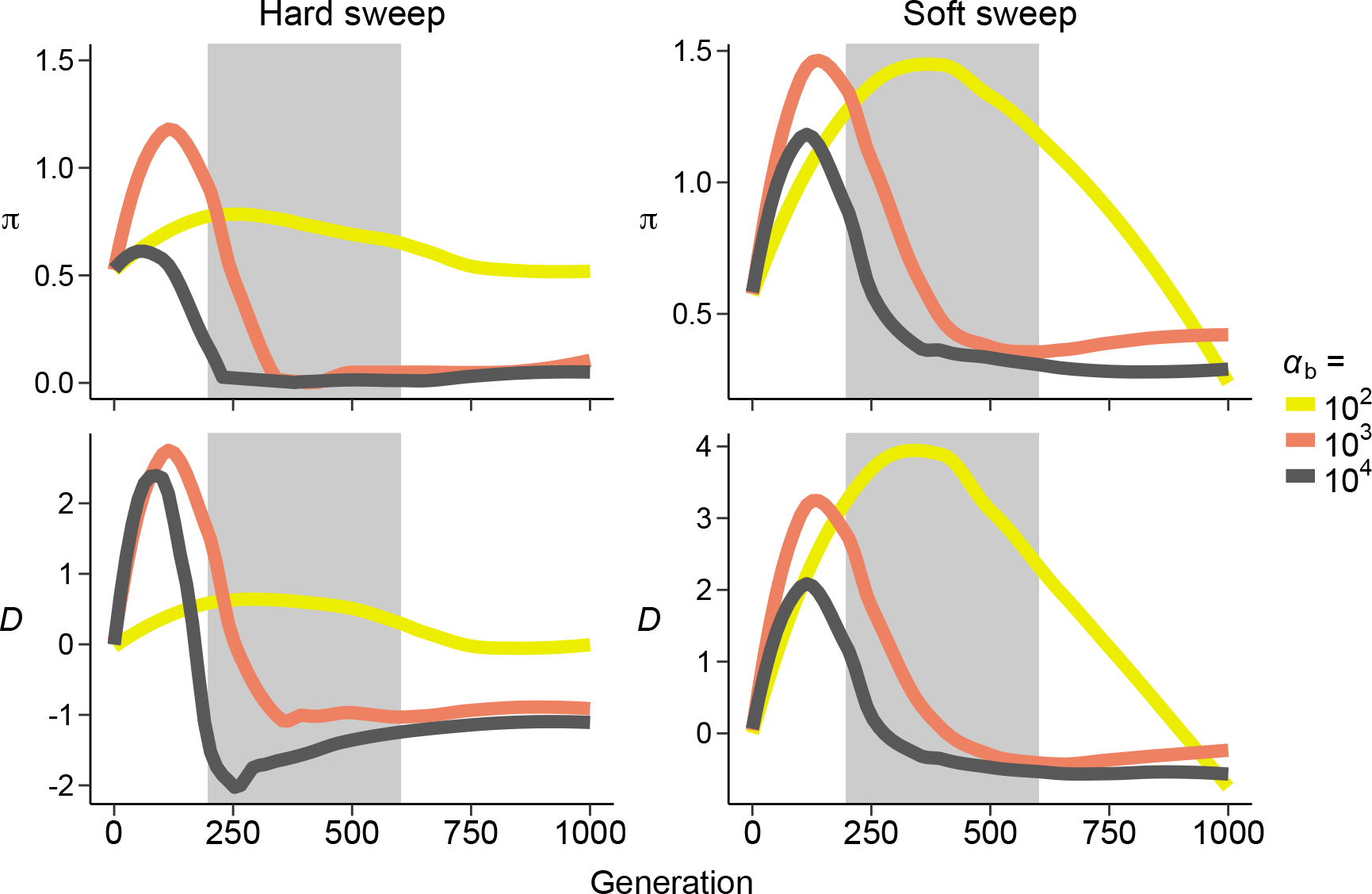
Nucleotide diversity *π* (×10^−3^) and Tajima’s *D* estimates simulated for 8 Kb area with parameters corresponding to T4 population. The simulations were ran as single-origin hard sweeps and as multiple-origin soft sweeps. For the latter case, we assumed an initial frequency of 0.05 for the adaptive allele. Shown are median estimates from 500 simulations. Shaded area marks the 95% confidence intervals for the estimated divergence time between T1 and T4 populations.

**Figure S12.**
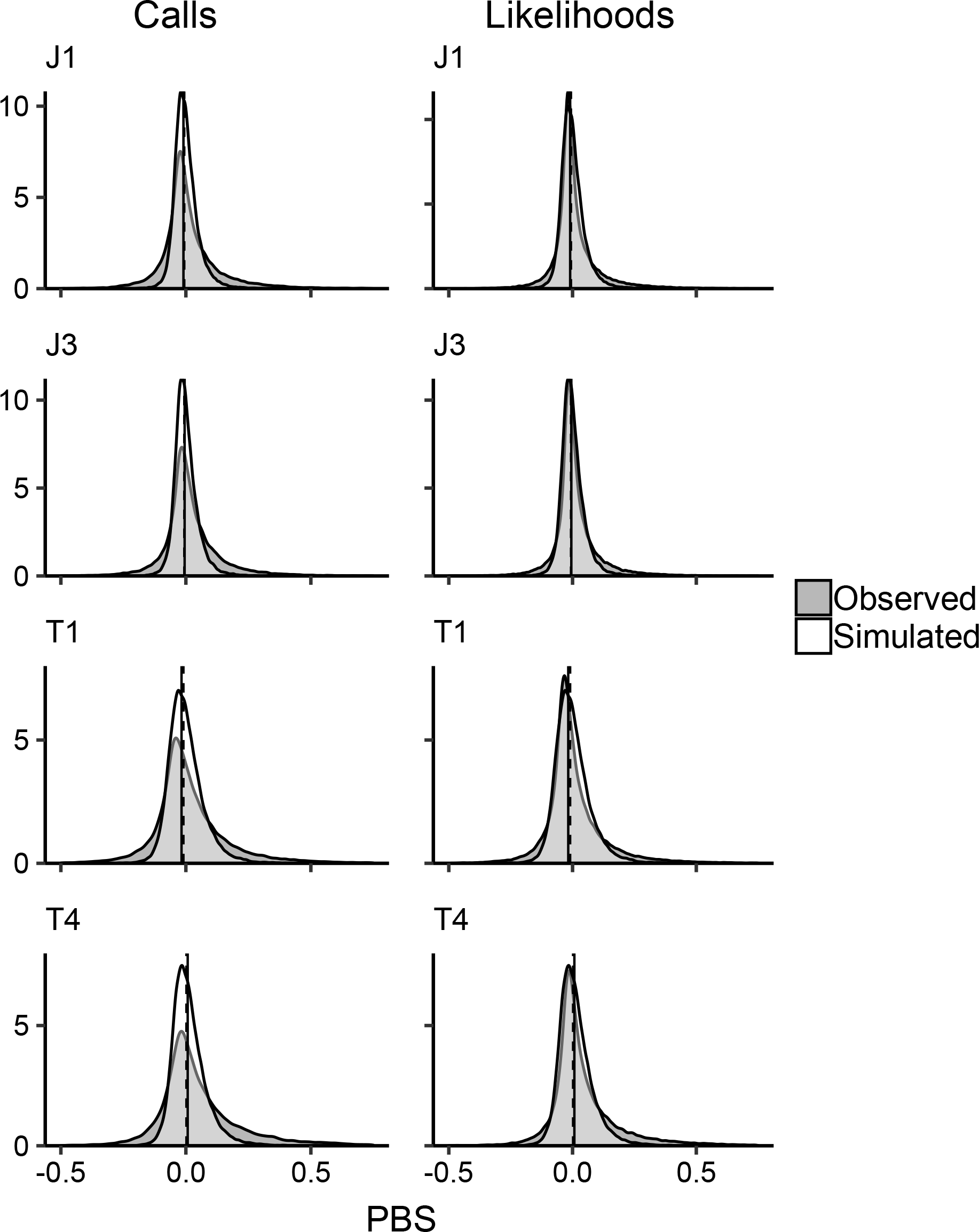
PBS distributions generated with SNP calls and genotype likelihoods compared against simulated neutral samples. Data were filtered using the following settings: mapping quality 30, site quality 20, genotype quality 20 (SNP calls only), minimum coverage 4×. SNPs were called with Freebayes and the genotype likelihoods were estimated with the GATK model in ANGSD.

**Figure S13.**
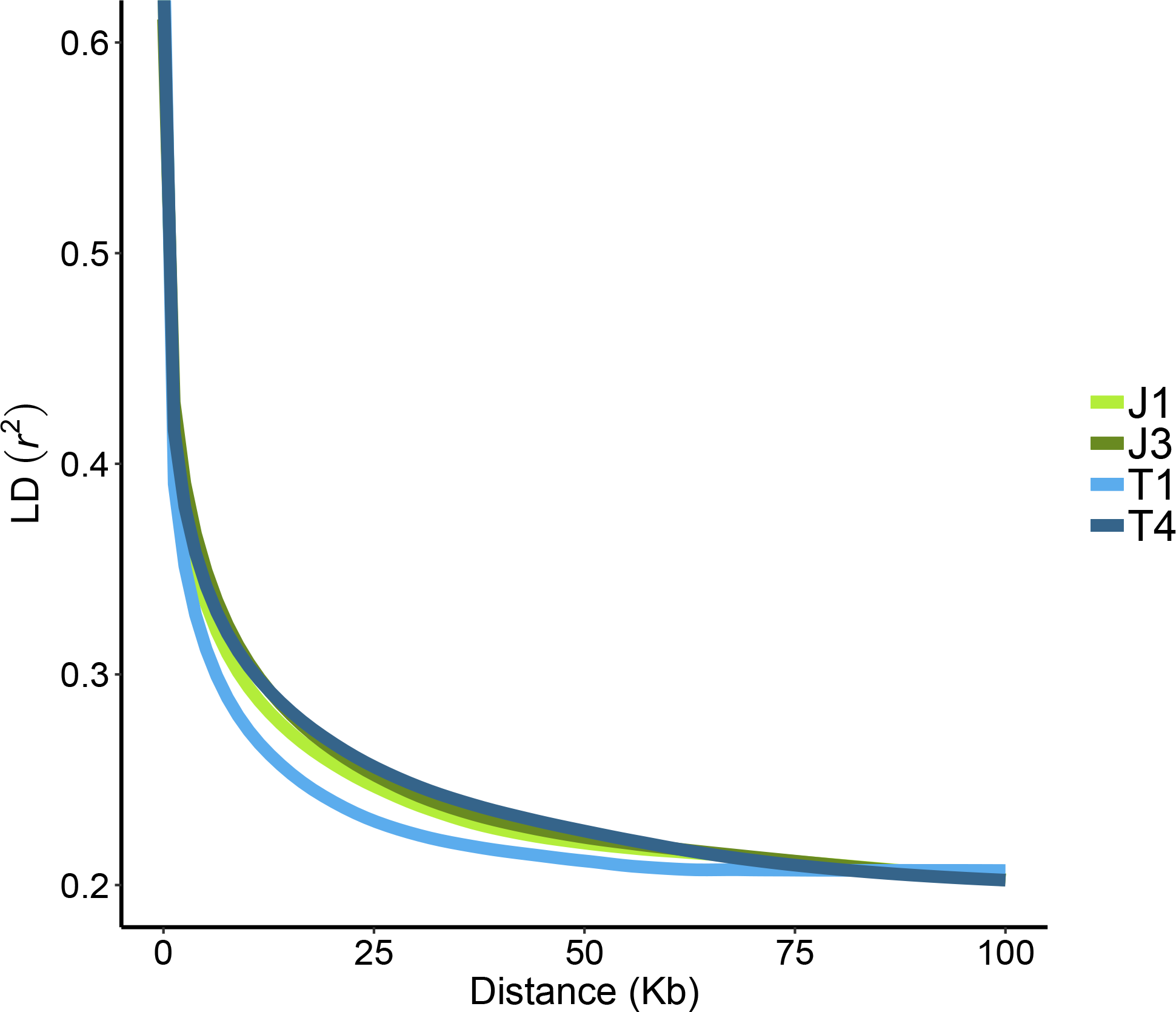
The decay of linkage disequilibrium (LD), as measured by *r*^2^ between SNP allele counts, with physical distance.

**Table S1.** Location information for the study populations.

| Population | Location | Lat (N) | Lon (E) | Altitude (m.a.s.l) |
| --- | --- | --- | --- | --- |
| J1 | Lom, Jotunheimen | 61°84′ | 8°57′ | 300 |
| J3 | Spiterstulen, Jotunheimen | 61°62′ | 8°40′ | 1100 |
| T1 | Sunndalsøra, Trollheimen | 62°66′ | 8°62′ | 10 |
| T4 | Nedre Kamtjern, Trollheimen | 62°74′ | 9°30′ | 1360 |
| SWE | Stubbsand, Sweden | 63°12′ | 18°57′ | 5 |
| GER | Plech, Germany | 49°39′ | 11°29′ | 400 |

**Table S2.** Pairwise *F*_ST_ estimates for the study populations.

| Population | J1 | J3 | T1 | T4 | SWE |
| --- | --- | --- | --- | --- | --- |
| J3 | 0.086 |  |  |  |  |
| T1 | 0.267 | 0.276 |  |  |  |
| T4 | 0.305 | 0.312 | 0.165 |  |  |
| SWE | 0.335 | 0.340 | 0.313 | 0.352 |  |
| GER | 0.412 | 0.425 | 0.369 | 0.413 | 0.417 |

Values are weighted genome-wide averages estimated for synonymous sites.

**Table S3.** The maximum likelihood function (*L*), Akaike information criterion (AIC) and Akaike weights (*w*) for each tested migration model. The best fitting models for each population comparison are in bold.

| Population model | Migration model | $\ln(L)$ | AIC | $w$ |
| --- | --- | --- | --- | --- |
| J1-J3-GER | No migration | -812,518.216 | 1,625,050 | $3.48 \times 10^{-19}$ |
| | From J1 to J3 | -812,517.500 | 1,625,051 | $2.12 \times 10^{-19}$ |
| | From J3 to J1 | -812,514.302 | 1,625,045 | $4.24 \times 10^{-18}$ |
|  | **Bidirectional migration** | **-812,473.580** | **1,624,965** | **1** |
|  | Bottleneck | -817,705.393 | 1,635,433 | 0 |
| T1-T4-GER | No migration | -794,034.865 | 1,588,084 | $1.03 \times 10^{-10}$ |
| | From T1 to T4 | -794,028.514 | 1,588,073 | $2.51 \times 10^{-8}$ |
| | From T4 to T1 | -794,017.947 | 1,588,052 | $9.11 \times 10^{-3}$ |
|  | **Bidirectional migration** | **-794,010.067** | **1,588,038** | **1** |
| | Bottleneck | -794,035.752 | 1,588,098 | $9.35 \times 10^{-14}$ |
| J1-T1-GER | No migration | -819,066.549 | 1,638,147 | $1.26 \times 10^{-15}$ |
|  | **From J1 to T1** | **-819,031.510** | **1,638,079** | **0.73** |
|  | From T1 to J1 | -819,032.371 | 1,638,081 | 0.27 |
| | Bidirectional migration | -819,059.187 | 1,638,136 | $3.07 \times 10^{-13}$ |
|  | Bottleneck | -820,688.170 | 1,641,398 | 0 |
| J3-T4-GER | **No migration** | **-864,472.383** | **1,728,959** | **1** |
| | From J3 to T4 | -864,495.367 | 1,729,007 | $3.78 \times 10^{-11}$ |
| | From T4 to J3 | -864,488.217 | 1,728,992 | $6.83 \times 10^{-8}$ |
| | Bidirectional migration | -864,519.009 | 1,729,056 | $8.64 \times 10^{-22}$ |
| | Bottleneck | -864,503.865 | 1,729,034 | $5.18 \times 10^{-17}$ |

**Table S4.**
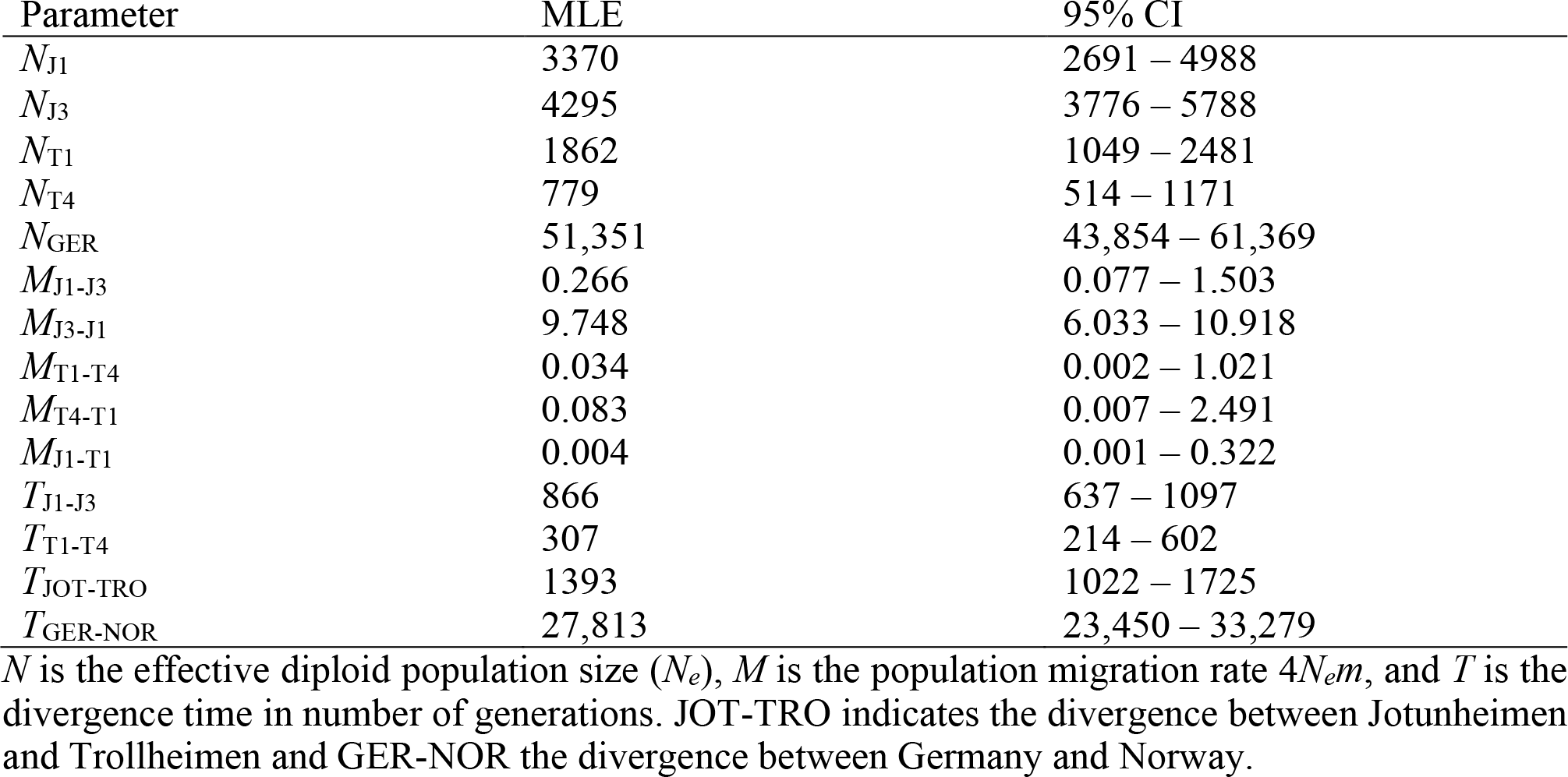
Maximum likelihood estimates (MLE) and their 95% confidence intervals for the demography parameters estimated from the observed data.

| Parameter | MLE | 95% CI |
| --- | --- | --- |
| $N_{J1}$ | 3370 | 2691 – 4988 |
| $N_{J3}$ | 4295 | 3776 – 5788 |
| $N_{T1}$ | 1862 | 1049 – 2481 |
| $N_{T4}$ | 779 | 514 – 1171 |
| $N_{GER}$ | 51,351 | 43,854 – 61,369 |
| $M_{J1-J3}$ | 0.266 | 0.077 – 1.503 |
| $M_{J3-J1}$ | 9.748 | 6.033 – 10.918 |
| $M_{T1-T4}$ | 0.034 | 0.002 – 1.021 |
| $M_{T4-T1}$ | 0.083 | 0.007 – 2.491 |
| $M_{J1-T1}$ | 0.004 | 0.001 – 0.322 |
| $T_{J1-J3}$ | 866 | 637 – 1097 |
| $T_{T1-T4}$ | 307 | 214 – 602 |
| $T_{JOT-TRO}$ | 1393 | 1022 – 1725 |
| $T_{GER-NOR}$ | 27,813 | 23,450 – 33,279 |

$N$ is the effective diploid population size ( $N_e$ ), $M$ is the population migration rate $4N_em$ , and $T$ is the divergence time in number of generations. JOT-TRO indicates the divergence between Jotunheimen and Trollheimen and GER-NOR the divergence between Germany and Norway.

**Table S5.**
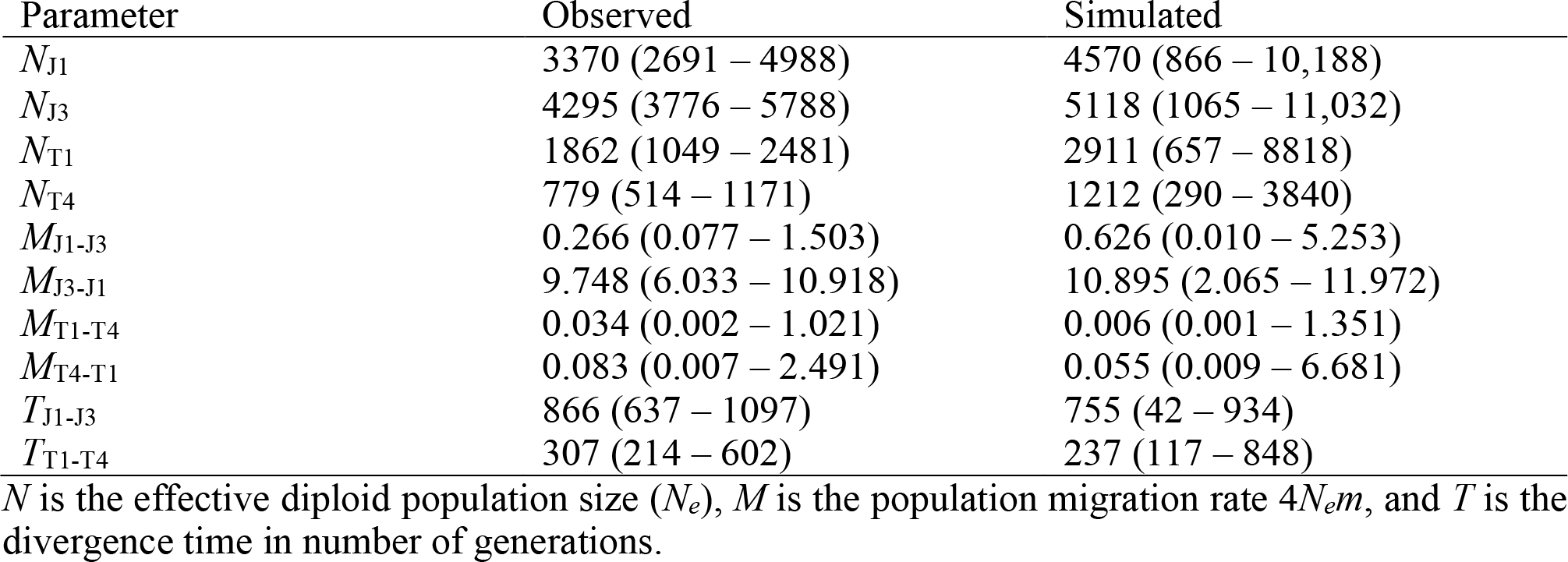
Demography estimates compared between observed and simulated data. Shown are maximum likelihood estimates and their 95% confidence intervals.

